# The cryo-EM-delineated mechanism underlying mimicry of CXCR4 agonism enables widespread stem cell neuroprotection in a mouse model of ALS

**DOI:** 10.1101/2025.07.08.663251

**Authors:** Xiaohong Sang, Haizhan Jiao, Qian Meng, Xiong Fang, Kartik S. Sundaram, Jiao Zhou, Yan Xu, Asuka I. W. Alvarado, Ruslan L. Nuryyev, Jitka Ourenik, Vaclav Ourednik, Iris S. Huang, Xiang Liu, Yuheng Mei, Tingli Qian, Aaron Ciechanover, Donald P. Pizzo, Michael A. Lane, Lyandysha V. Zholudeva, Jing An, Evan Y. Snyder, Hongli Hu, Ziwei Huang

**Affiliations:** Ciechanover Institute of Precision and Regenerative Medicine, School of Medicine, The Chinese University of Hong Kong, Shenzhen 518172, China; Kobilka Institute of Innovative Drug Discovery, School of Medicine, The Chinese University of Hong Kong, Shenzhen 518172, China; School of Life Sciences, Tsinghua University, Beijing 100084, China; Center for Stem Cells and Regenerative Medicine, Sanford Burnham Prebys Medical Discovery Institute, La Jolla, CA 92037, USA; Division of Infectious Diseases and Global Public Health, Department of Medicine, School of Medicine, University of California San Diego, La Jolla, CA 92037, USA; Sanford Consortium for Regenerative Medicine, La Jolla, CA 92037, USA; School of Chemistry and Chemical Engineering, Guangdong Pharmaceutical University, Zhongshan 528458, China; Technion Rappaport Integrated Cancer Center, The Rappaport Faculty of Medicine and Research Institute, Technion-Israel Institute of Technology, Haifa 3109601, Israel; Department of Pathology, University of California San Diego, La Jolla, CA 92037, USA; Drexel University, Philadelphia, PA 19104, USA; Gladstone Institutes, San Francisco, CA 94158, USA

## Abstract

G-protein coupled receptors (GPCRs) are transmembrane proteins that mediate a range of signaling functions and, therefore, offer targets for a number of therapeutic interventions. Chemokine receptor CXCR4, a GPCR, plays versatile roles in normal and abnormal physiological processes. Synthetic CXCR4 antagonists have been extensively studied and approved for the clinical treatment of cancer and other diseases. We recently elucidated the structural mechanisms underlying CXCR4 antagonism using cryogenic electron microscopy (cryo-EM). CXCR4 agonism by synthetic molecules is an unanticipated therapeutic intervention we recently unveiled. The structural mechanisms underlying those actions remain poorly understood yet could help elucidate a new class of drugs. Here we demonstrate a synthetic dual-moiety strategy that combines simplified agonistic and antagonistic moieties taken from natural agonistic and antagonistic chemokines, respectively, to design de novo peptide mimics of biological function of natural CXCR4 agonist SDF-1α. Two peptides so generated, SDV1a and SDVX1 were shown to mimic the action of SDF-1α in activating CXCR4 signaling pathways and cell migration. The structural mechanism of these peptides in the mimicry of CXCR4 agonism was illustrated by cryo-EM structures of CXCR4 bound and activated by the peptides in the presence of G protein, revealing common interactions with the receptor by these peptides in comparison with SDF-1α that explain their close mimicry and conformational changes leading to CXCR4 signal activation. The therapeutic benefit of one of these peptides, SDV1a, was demonstrated in the SOD1^G93A^ mouse model of the spinal motor neuron degenerative disease, amyotrophic lateral sclerosis (ALS) wherein the success of neuroprotective actions of transplanted human neural stem cells (hNSCs) is directly correlated with the expanse of diseased neuroaxis traversed by the donor cells; SDV1a enabled broader neuroprotective coverage while also permitting a much less invasive route of cell administration for extending life. Taken together, these results provide insights into the structural determinants of therapeutic CXCR4 agonism which may allow the design of adjunctive drugs that improve cell-based treatments of central nervous system (CNS) diseases.

## Main

The G-protein coupled receptors (GPCRs) are a superfamily of membrane proteins that mediate cellular signaling critical for a wide range of biological processes ^1^ and, hence, are the targets for many drugs for a variety of diseases ^2^. The functions of GPCRs in human physiology and pathology can be manipulated and studied by ligands of these receptors, some represented in the current clinical arsenal ^3^. There have been many studies to understand the structures and function of GPCRs and explore their clinical potentials. Nevertheless, many questions remain regarding the structural mechanisms of GPCR signaling.

The chemokine receptor CXCR4 is a GPCR implicated in various physiological and pathological processes such as HIV entry ^4,5^, cancer metastasis ^6^, inflammation ^7^ and stem cell trafficking ^8^. Most drug discovery efforts have focused on CXCR4 antagonism which appeared to have the greatest clinical utility. Such antagonism has included peptides, small molecules and nanobodies ^9,10^, among them two small molecule antagonist drugs approved for clinical uses, AMD3100 ^11^ and AMD070 ^12^. The structural mechanisms of some of these antagonists bound to CXCR4 receptor have been determined through X-ray crystallography, e.g., for the peptide antagonist CVX15 ^13^, the small molecule antagonist It1t ^13^, and the viral chemokine vMIP-II ^14^. Of late, our laboratories have applied cryo-electron microscopy (cryo-EM) technology to solve the CXCR4 bound structures of the two clinical antagonist drugs AMD3100 and AMD070 and a small molecule antagonist HF51116 ^15^.

Previously, we introduced the notion that some aspects of CXCR4 agonism might have therapeutic benefit ^16^ – for example in the guidance of reparative cells to regions of central nervous system (CNS) pathology and the induction of intracellular neural repair pathways. Maximizing those actions, while eliminating or minimizing CXCR4’s undesirable downstream signaling (e.g., inflammation, apoptosis) requires exquisite dissection of CXCR4’s signaling determinants to design agonist ligands with controllably selective actions. Compared to CXCR4 antagonists, the therapeutic development of CXCR4 agonists is less explored; structural understanding of synthetic CXCR4 agonism poorly understood, but exceptionally translationally important.

In this study, we applied cryo-EM to determine the structures of CXCR4-G protein complexes bound and activated by two de novo designed and chemically synthesized agonist peptides, SDV1a and SDVX1. These novel agonists were tested in a panel of cellular biological assays and shown to mimic the biological activity of natural wild-type agonist ligand SDF-1α in activating CXCR4-G protein complex downstream signaling pathways and promoting consequent cell migration. Such functional mimicry of the natural chemokine ligand with much smaller and simplified synthetic molecules is explained and justified by the similar impact on CXCR4-G protein complex structures inflicted by these synthetic mimics versus the wild-type ligand as revealed in the cryo-EM structures. The translational utility of these peptides was supported by using one of them in the treatment of the SOD1^G93A^ mouse model of the spinal motor neuron (MN) degenerative disease, amyotrophic lateral sclerosis (ALS) ^17^ wherein the success of neuroprotective actions of transplanted human neural stem cells (hNSCs) is directly correlated with the expanse of diseased neuroaxis traversed by the donor cells. When hNSCs and SDV1a were co-administered using a novel minimally-invasive protocol (enabled by SDV1a), the treated ALS mice, which typically die at ∼150 days-of-life, lived at least 3-times as long, meeting functional criteria and with evidence of preserved MNs, until the experiment was artificially terminated. In other words, the peptide enabled greater neuroprotection while reducing the invasiveness of cell administration. Taken together, these structural and animal studies validate the design and translational potential of our synthetic GPCR agonist strategy of linking two simplified moieties – one responsible for pure GPCR signaling and the other for pure binding – to create a de novo drug lead that mimics the desirable aspects of CXCR4 signaling and might be used to enhance the efficacy of cell-based treatments of CNS diseases.

### De novo design and biological characterization of synthetic CXCR4 agonists

To dissert and study the structural and functional determinants of CXCR4 agonism, we designed a synthetic probe that separates the receptor’s two functions: signaling and binding, as we previously described ^18^. For the signaling moiety, we truncated the 68 amino-acid (AA) N-terminal signaling motif of CXCR4’s natural ligand SDF-1α to just 8 AAs. Into SDF-1α’s binding motif, we swapped in the residues of CXCR4’s strong antagonist, viral vMIP-II. This new traceable synthetic “dual moiety” peptide we called “SDV1a”. We then further reduced the chemical structure of SDV1a to its even more minimalistic basic functional elements, and called that peptide “SDVX1” (Fig. 1A, Table 1, and Extended Data Fig. 1). For SDVX1, we retained the critical signaling moiety of SDV1a (i.e., its N-terminal residues 1-8), but shortened the C-terminal binding moiety of SDV1a from 21 to only 10 residues. Surprisingly, although much smaller than SDV1a, SDVX1 showed enhanced CXCR4 binding affinity with IC_50_ of 760 nM (Fig. 1B and Table 1). Furthermore, while both SDVX1 and SDV1a promoted cell migration in an “inverted parabola” dose-dependent manner, the optimal concentration for SDVX1 at 0.5 μM showed it to be 10-fold more potent than SDV1a (Fig. 1C and Table 1). Note that the design of SDVX1 contains NalRR (i.e., Nal-Arg-Arg) at its C-terminus, having been extracted from the N-terminus of a CXCR4 peptide antagonist CVX15 ^13^. Furthermore, SDV1a and SDVX1 are quite specific in their binding of CXCR4; for example, they do not bind a closely-related chemokine receptor, CCR5, even at very high concentrations (50 μM and 10 μM, respectively) (Extended Data Fig. 2A).

**Figure 1.**
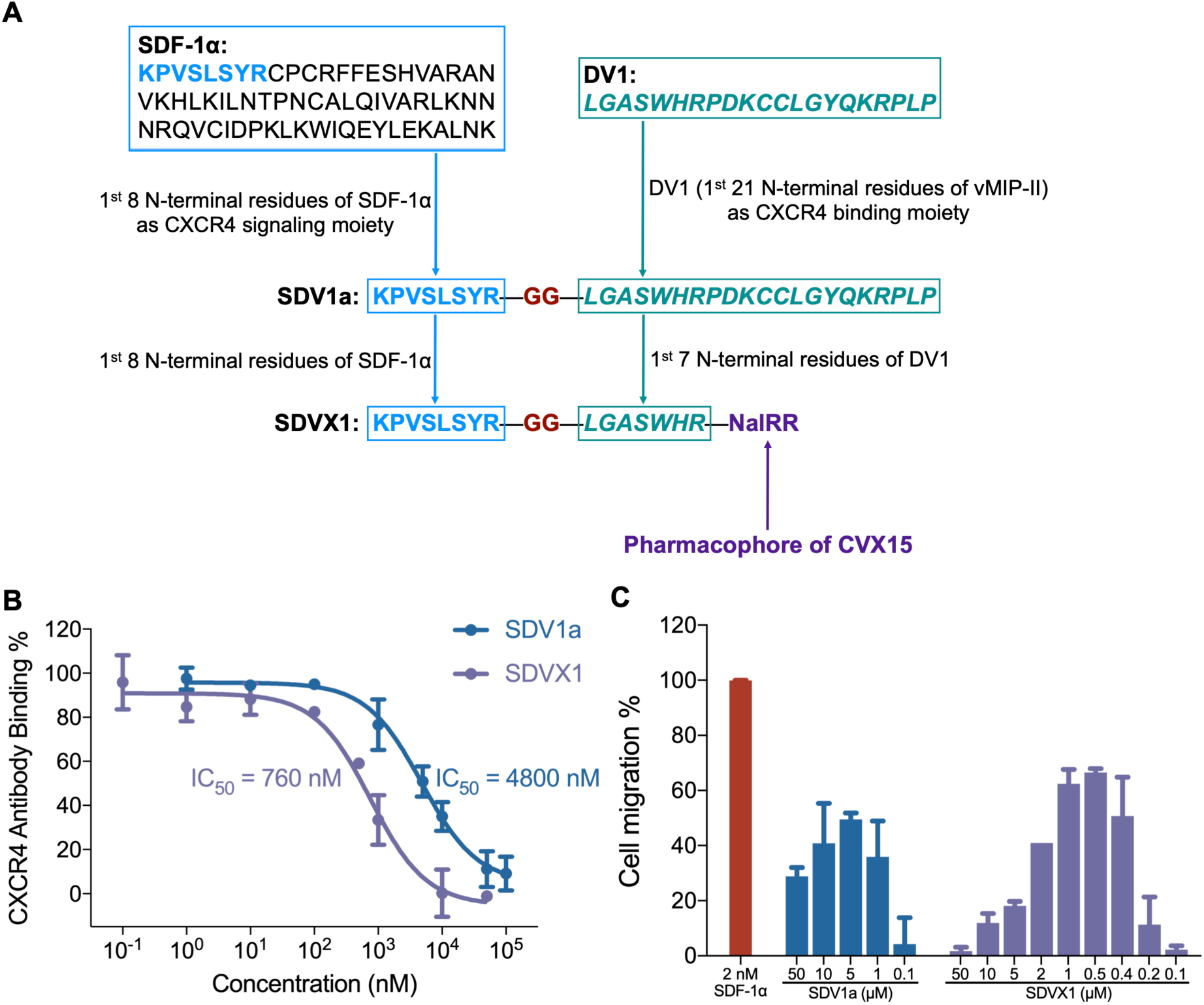
Design and characterization of SDV1a and SDVX1. **(A)** The de novo design of SDV1a and SDVX1 using a dual-moiety strategy. For SDV1a, the peptide fragment KPVSLSYR from agonist chemokine SDF-1α N-terminus was selected as the CXCR4 signaling moiety (in blue) whereas DV1 sequence (in dark teal) corresponding to the 21 N-terminal residues of antagonist chemokine vMIP-II served as the CXCR4 binding moiety chemically linked to the above described signaling moiety via two glycine residues (in dark red), thus creating a simplified hybrid molecule yet with both CXCR4 binding and activation functions like the natural ligand SDF-1α. For SDVX1, while the signaling moiety and linker stayed the same, the DV1 binding moiety was reduced significantly to only 10 residues, *LGASWHR*, added with another antagonist peptide CVX15-truncated short fragment NalRR (in dark purple). **(B)** The binding affinity of SDV1a and SDVX1 on CXCR4. **(C)** The cell migration-promoting activity of SDV1a and SDVX1. 100% migration activity was promoted by SDF-1α (2 nM). These results demonstrate the functional mimicry of SDF-1α by these synthetic agonist peptides in promoting CXCR4-mediated cell migration.

**Table 1.** Sequences and activities of two CXCR4 de novo agonist peptides.

| Peptide name | Sequence<sup>a</sup> | CXCR4 binding potency<br>(IC<sub>50</sub>) | Cell migration<sup>b</sup> |
| --- | --- | --- | --- |
| SDV1a | KPVSLSYR-GG-<u>LGASWHRPDKCCLGYQKRPLP</u> | 4800 nM | ~50% (5 µM) |
| SDVX1 | KPVSLSYR-GG-<u>LGASWHR</u>-NaIRR | 760 nM | ~70% (0.5 µM) |

a. The D-amino acids are underlined and shown in *italics*.
b. 100% migration activity is promoted by SDF-1α (2 nM).

We analyzed further the agonistic effect of SDV1a and SDVX1 on CXCR4 downstream signaling pathways. When activated by its natural ligand SDF-1α, CXCR4 triggers downstream G protein-dependent signaling pathways, leading to decrease of cellular cAMP, increase of calcium in the cytoplasm, and activation of MAPK/ERK and RhoA signaling pathways ^19^. We employed a luciferase reporter system to assess the activation of these signaling pathways ^20^. This system includes the cAMP response element (CRE) for detecting cAMP levels, the nuclear factor of activated T-cells response element (NFAT-RE) for activation of intracellular calcium pathway, the serum response element (SRE) for activation of MAPK/ERK signaling pathway, and the serum response factor response element (SRF-RE) for activation of RhoA signaling pathway. In a dose-dependent manner, both SDV1a and SDVX1 could exerted effects similar to SDF-1α’s on cAMP decrease, calcium increase, and activation of MAPK/ERK, and RhoA (Fig. 2A-D), thus demonstrating the functional mimicry of the natural agonist ligand by these synthetic molecules.

**Figure 2.**
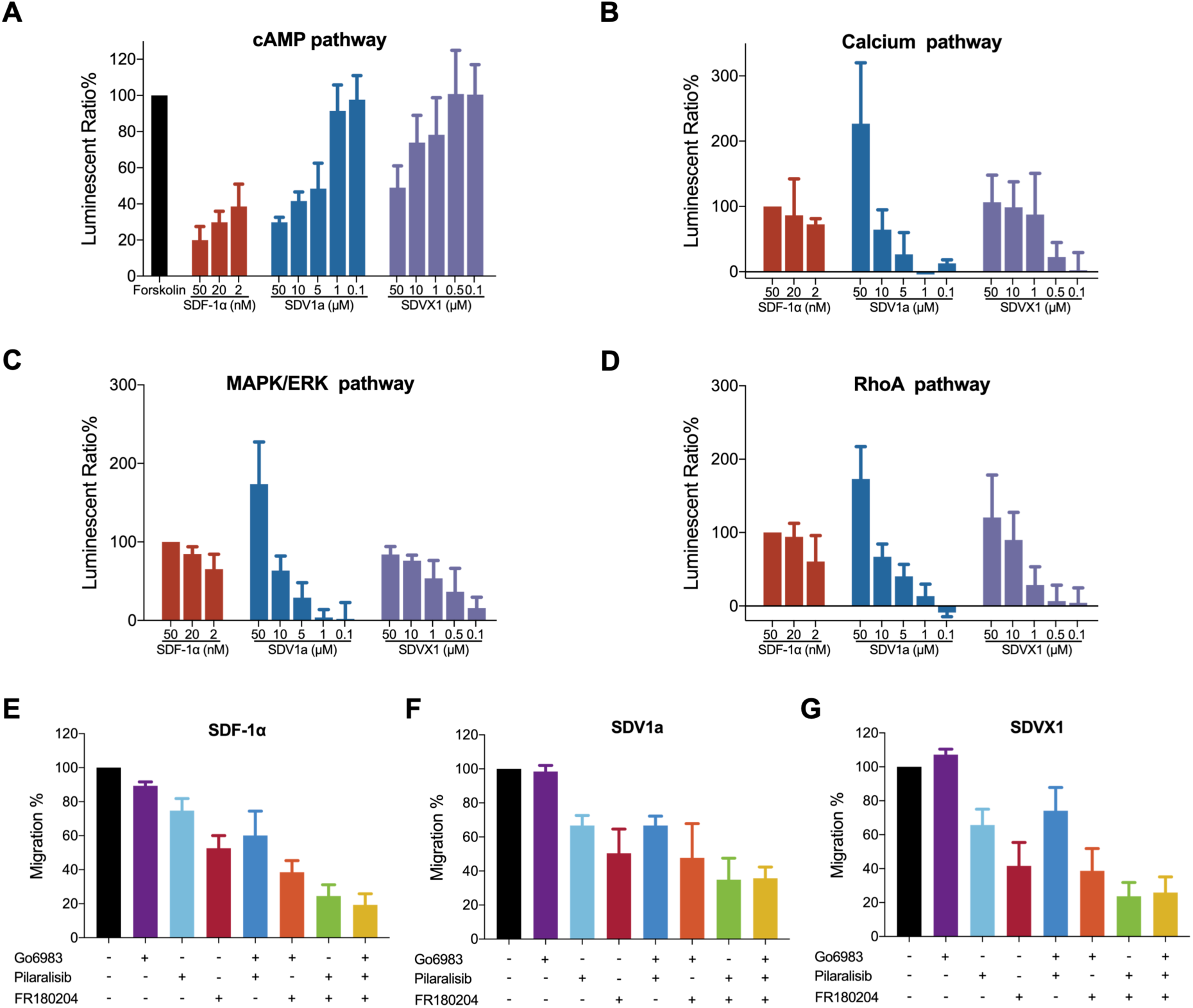
The effects of SDV1a and SDVX1 similar to SDF-1α on CXCR4 signaling pathways. **(A)** The agonistic activity of SDF-1α, SDV1a and SDVX1 on cAMP signaling pathway. Forskolin (2.5 μg/mL) was used to stimulate CHO cells to increase the production of intracellular cAMP. **(B)** The agonistic activity of SDF-1α, SDV1a and SDVX1 on calcium signaling pathway. **(C)** The agonistic activity of SDF-1α, SDV1a and SDVX1 on MAPK/ERK signaling pathway. **(D)** The agonistic activity of SDF-1α, SDV1a and SDVX1 on RhoA signaling pathway. **(E)** Effect of signaling pathway inhibitors on cell migration induced by SDF-1α. **(F)** Effect of signaling pathway inhibitors on cell migration induced by SDV1a. **(G)** Effect of signaling pathway inhibitors on cell migration induced by SDVX1. 20 μM of Go6983 (PKC inhibitor), 20 μM of Pilaralisib (PI3K inhibitor), and 100 μM of FR180204 (ERK1/2 inhibitor) were used for signaling pathway inhibitors in E-G. Together with the data shown in Fig. 1, these results further illustrate the biological activities of these two synthetic peptides in mimicking the effects of SDF-1α on CXCR4 functions.

We also observed a diminished ability of SDV1a and SDVX1 to promote cell migration by selectively blocking 3 signaling pathways involved in cell migration, indicating their importance for the peptides’ chemoattraction properties: Go6983 to block PKC, Pilaralisib to block PI3K, FR180204 to block ERK1/2. Treatment with each inhibitor alone inhibited cell migration to a certain extent, with co-treatments of two or three inhibiting synergistically. CXCR4’s natural ligand, SDF-1α, was used a baseline (Fig. 2E). The cell migration induced by SDV1a and SDVX1 was inhibited by Pilaralisib and FR180204, but, in contrast to SDF-1α, not Go6983, and no synergism was observed when Go6983 was combined with Pilaralisib and/or FR180204 (Fig. 2F and 2G). These results suggest that the migration promoting mechanism of these synthetic agonists, while sharing some signaling pathways with SDF-1α, also employs additional parallel pathways that make them potentially better agents for directing the migration of cells.

### Cryo-EM structures of CXCR4-Gi complexes bound and activated by synthetic agonists

We performed cryo-EM to understand the structural mechanism of CXCR4 agonism activated by SDV1a and SDVX1. We co-expressed and purified the CXCR4-Gi complex (Extended Data Fig. 2B). For SDV1a, the CXCR4-Gi-scFv16-SDV1a complex was obtained by adding GPCR/G-protein complex-stabilizing antibody scFv16 and further purified by affinity chromatography (Extended Data Fig. 2C). The structure of CXCR4-Gi-scFv16-SDV1a complex was subsequently resolved using cryo-EM single particle analysis (Extended Data Fig. 3A and Extended Data Table 1), achieving a density map with an overall resolution of 2.8 Å (Fig. 3A, left). An atomic model was built based on the resolved structure (Fig. 3A, right). Clear densities for the seven transmembrane helices were observed (Extended Data Fig. 3B), along with a density in the orthosteric binding pocket of CXCR4 which appeared to accommodate the N-terminal eight residues of SDV1a (Fig. 3A, middle). The densities corresponding to the C-terminal residues of SDV1a were unresolved due to density blurring likely attributed to the flexibility in this region which is consistent with unresolved C-terminal structure of natural SDF-1α bound to CXCR4 reported recently ^21^.

**Figure 3.**
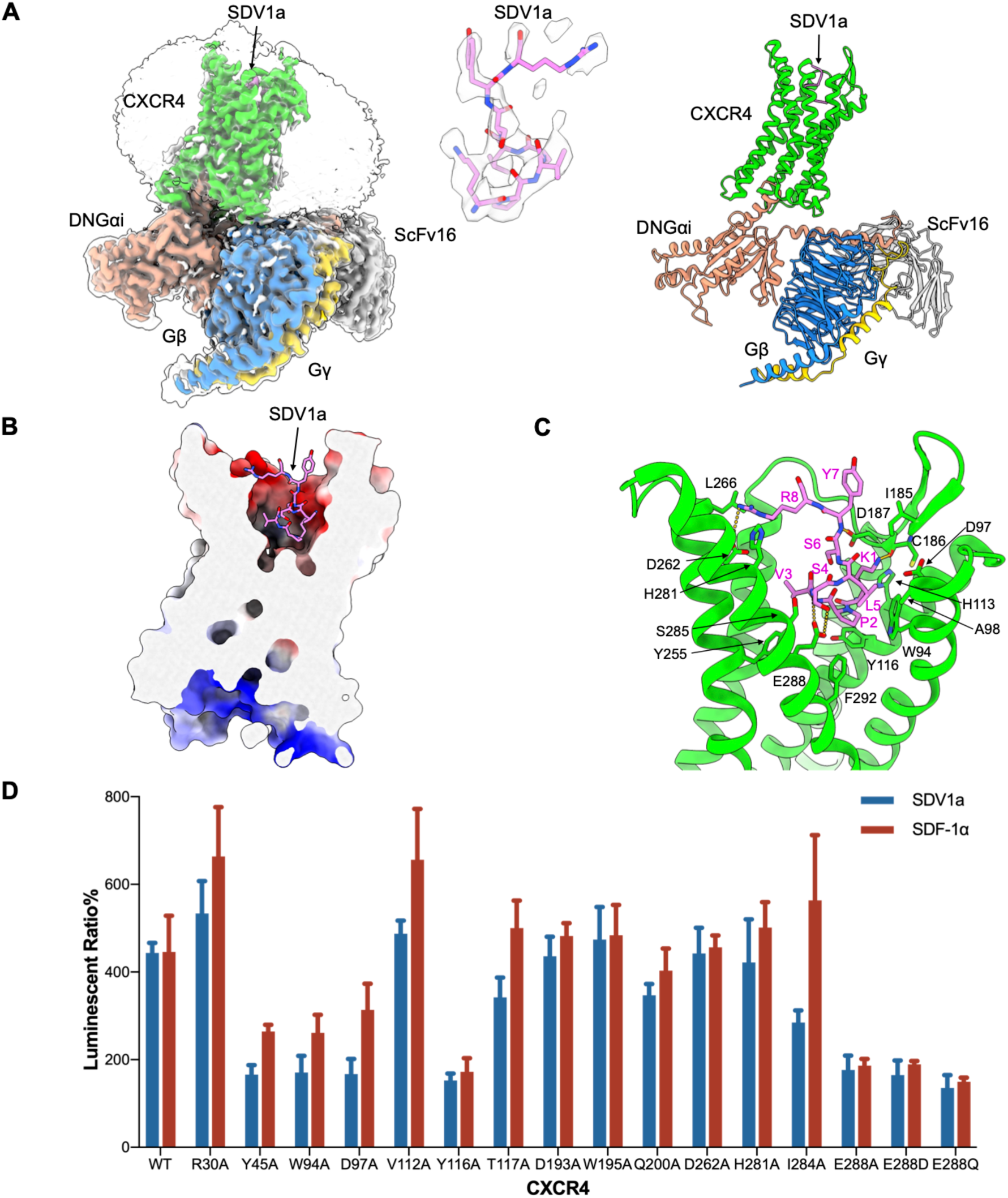
Cryo-EM structure of CXCR4-SDV1a and functional validation of the structural information by site directed mutagenesis. **(A)** The cryo-EM density (left) and the atomic model (right) of the SDV1a-CXCR4-DNGi-scFv16 complex. SDV1a, CXCR4, DNGαi, Gβ, Gγ, and scFv16 were colored violet, green, salmon, dodger blue, gold, and gray, respectively. The local density of the N-terminal eight residues of SDV1a is presented as well. **(B)** The binding pocket of SDV1a in CXCR4. CXCR4 is shown as a surfaced model and colored by electronic potential, while the N-terminus of SDV1a is shown as sticks and colored violet. **(C)** Interaction between SDV1a and CXCR4. CXCR4 is shown as a cartoon and colored green, and residues involved in interaction are shown as sticks. The N-terminus of SDV1a is shown as sticks and colored violet. Hydrogen bonds are indicated by yellow dotted lines. **(D)** Effects of CXCR4 mutations on signaling activated by SDV1a and SDF-1α. These site directed mutagenesis results support the important functional roles of CXCR4 residues as revealed by the cryo-EM structures.

The N-terminal 8 residues of SDV1a are positioned within the negatively charged central pocket of CXCR4 (Fig. 3B). Specifically, the side chain of Lys1^SDV1a^ establishes a hydrogen bond with the main chain of Cys186^ECL2^, further stabilized by hydrophobic stacking interactions with Ile185^ECL2^ and His113^3.29^ (Fig. 3C). Pro2^SDV1a^ engages in hydrophobic stacking with Trp94^2.60^, Tyr116^3.32^, and Phe292^7.43^ (Fig. 3C). Val3^SDV1a^ interacts hydrophobically with the side chain of Tyr255^6.51^ (Fig. 3C). Ser4^SDV1a^ is coordinated by Glu288^7.39^ through hydrogen bonds and interacts with Ser285^7.36^ via additional hydrophobic contacts (Fig. 3C). Leu5^SDV1a^ establishes hydrophobic interactions with Asp97^2.63^ and Ala98^2.64^ (Fig. 3C). The main chain of Tyr7^SDV1a^ forms a hydrogen bond with the side chain of Asp187^ECL2^, while the side chain of Arg8^SDV1a^ interacts with Asp262^6.58^ and engages in hydrophobic interactions with Leu266^6.62^ and His281^7.32^ (Fig. 3C).

Similar to SDV1a, we determined the cryo-EM structure of SDVX1 bound to CXCR4. The CXCR4-Gi-scFv16-SDVX1 complex was purified by affinity chromatography (Extended Data Fig. 2D). Single-particle analysis of the cryo-EM dataset yielded a density map with an overall resolution of 2.7 Å for this complex (Extended Data Fig. 4A and Extended Data Table 1). Notably, the densities of each transmembrane helix were clearly distinguished (Extended Data Fig. 4B). Similarly, the N-terminal 8 residues of SDVX1 are also positioned within the CXCR4 pocket (Fig. 4A and 4B), and the interaction between SDVX1 and CXCR4 closely resembles that of SDV1a with CXCR4 (Fig. 4C and 3C). Specifically, the side chain of Lys1^SDVX1^ had a hydrogen bond with the main chain of Cys186^ECL2^ and a salt bridge with Asp97^2.63^ (Fig. 4C). Pro2^SDVX1^ engaged in hydrophobic stacking with Trp94^2.60^, His113^3.29^, and Tyr116^3.32^ (Fig. 4C). Val3^SDVX1^ interacted hydrophobically with the side chain of Tyr255^6.51^ (Fig. 4C). Ser4^SDVX1^ was coordinated by Glu288^7.39^ through hydrogen bonds and interacted with Ser285^7.36^ via additional hydrophobic contacts (Fig. 4C). Leu5^SDVX1^ had hydrophobic interactions with Ala98^2.64^ (Fig. 4C). The main chain of Tyr7^SDVX1^ formed a hydrogen bond with the side chain of Asp187^ECL2^, while the side chain of Arg8^SDVX1^ had hydrogen bonds with Asp262^6.58^ and His281^7.32^ and salt bridges with Asp262^6.58^ (Fig. 4C).

**Figure 4.**
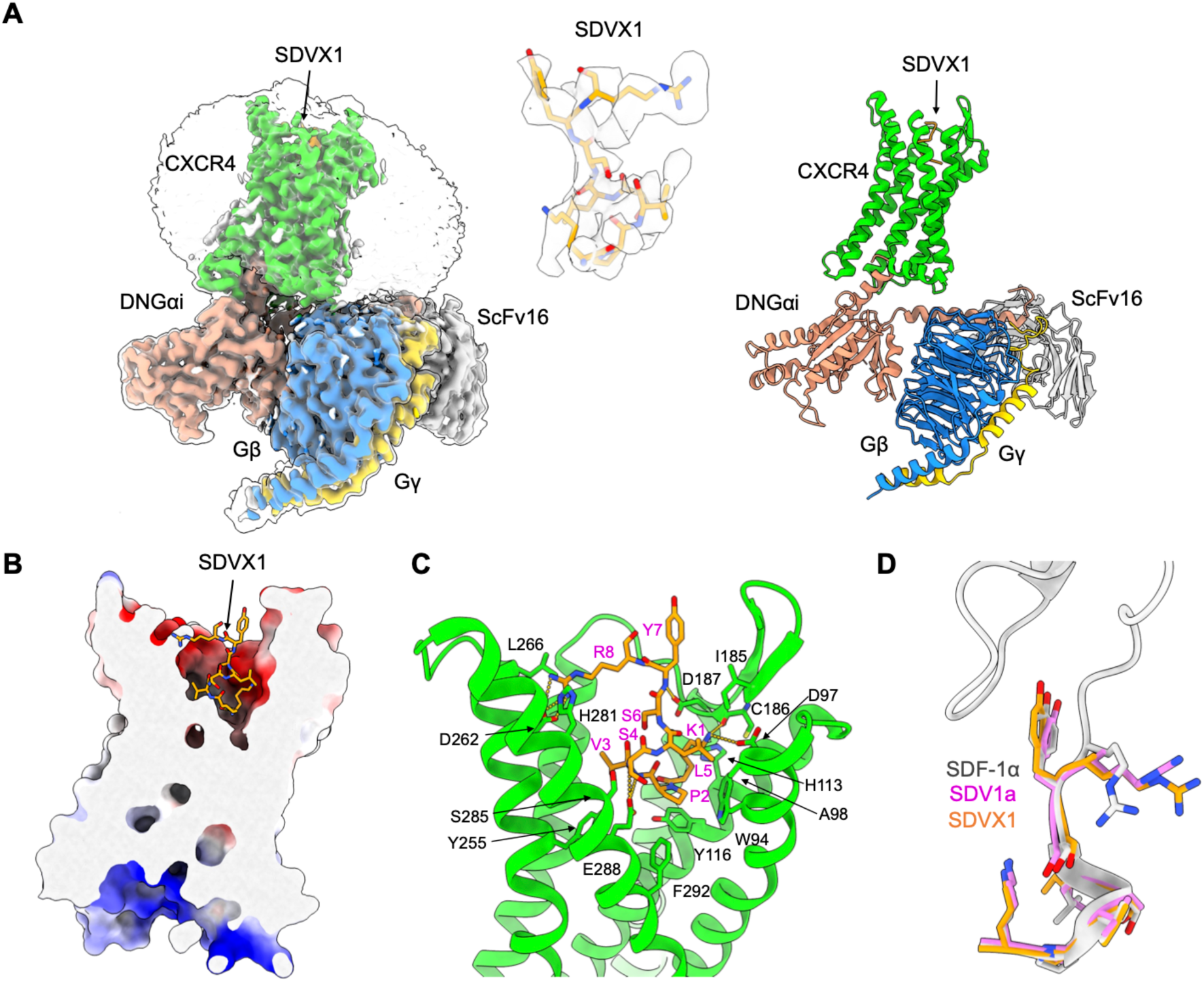
Cryo-EM structure of CXCR4-SDVX1. **(A)** The cryo-EM density (left) and the atomic model (right) of the SDVX1-CXCR4-DNGi-scFv16 complex. SDVX1, CXCR4, DNGαi, Gβ, Gγ, and scFv16 were colored orange, green, salmon, dodger blue, gold, and gray, respectively. The local density of the N-terminal eight residues of SDVX1 is presented as well. **(B)** The binding pocket of SDVX1 in CXCR4. CXCR4 is shown as surfaced model and colored by electronic potential, while the N-terminus of SDVX1 is shown as sticks and colored orange. **(C)** Interaction between SDVX1 and CXCR4. CXCR4 is shown as cartoon and colored green, and residues involved in interaction are shown as sticks. The N-terminus of SDVX1 is shown as sticks and colored orange. Hydrogen bonds are indicated by yellow dotted lines. **(D)** Superposition of SDF-1α and N-terminal eight residues of SDV1a and SDVX1, illustrating the close CXCR4 bound structural mimicry of SDF-1α by SDV1a and SDVX1 consistent with their functional mimicry in activating CXCR4 signaling.

The CXCR4 bound structure of the N-terminal residues 1-8 of SDVX1 was found to be very similar to the corresponding segments of SDV1a and SDF-1α bound to CXCR4 (Fig. 4D). A comparison of the N-termini of these two peptide agonists and SDF-1α in complex with CXCR4 reveals that the mainchain atoms of their N-terminal eight residues are positioned in closely aligned conformations (Fig. 4D). In contrast, while the sidechain positions of the first six residues (K1-S6) remain relatively constant, the sidechain interactions of residues Y7 and R8 exhibit increased variability (Fig. 4D). These observations suggest that the first six residues of SDVX1, SDV1a, and SDF-1α bind to CXCR4 in a stable and conserved manner, whereas the binding mode of the subsequent residues becomes more divergent. This implies that the unresolved regions of the peptides may adopt more dynamic binding modes without a fixed structure.

To further compare and analyze these structures, we utilized the residue-residue contact score (RRCS) analysis, an approach that quantifies the contact strength between residue pairs by summing all possible heavy-atom contact distances ^22^. By comparing the contact differences (ΔRRCS) of identical residue pairs between the two systems, we identified residues exhibiting significant structural changes. In total, ΔRRCS calculations were performed on 1,672 residue pairs in the CXCR4 systems bound by SDV1a and SDF-1α, revealing 21 residue pairs with |ΔRRCS| values exceeding 3 (Extended Data Table 2). Further structural analysis of these 21 residue pairs identified two distinct regions of significant structural perturbation: a TM1 cluster comprising residues N35^1.29^, F36^1.30^, N37^1.31^, K38^1.32^, I39^1.33^, F40^1.34^, and L41^1.35^, and a TM7 cluster formed by residues C274^7.25^, E275^7.26^, E277^7.28^, H281^7.32^, K282^7.33^, and W283^7.34^ (Fig. 5A). These findings highlight localized conformational rearrangements in the TM1 and TM7 regions that are induced by distinct CXCR4 agonists. We also conducted the ΔRRCS calculations between SDVX1 and SDF-1α systems, and similar TM1 and TM7 perturbations were identified.

**Figure 5.**
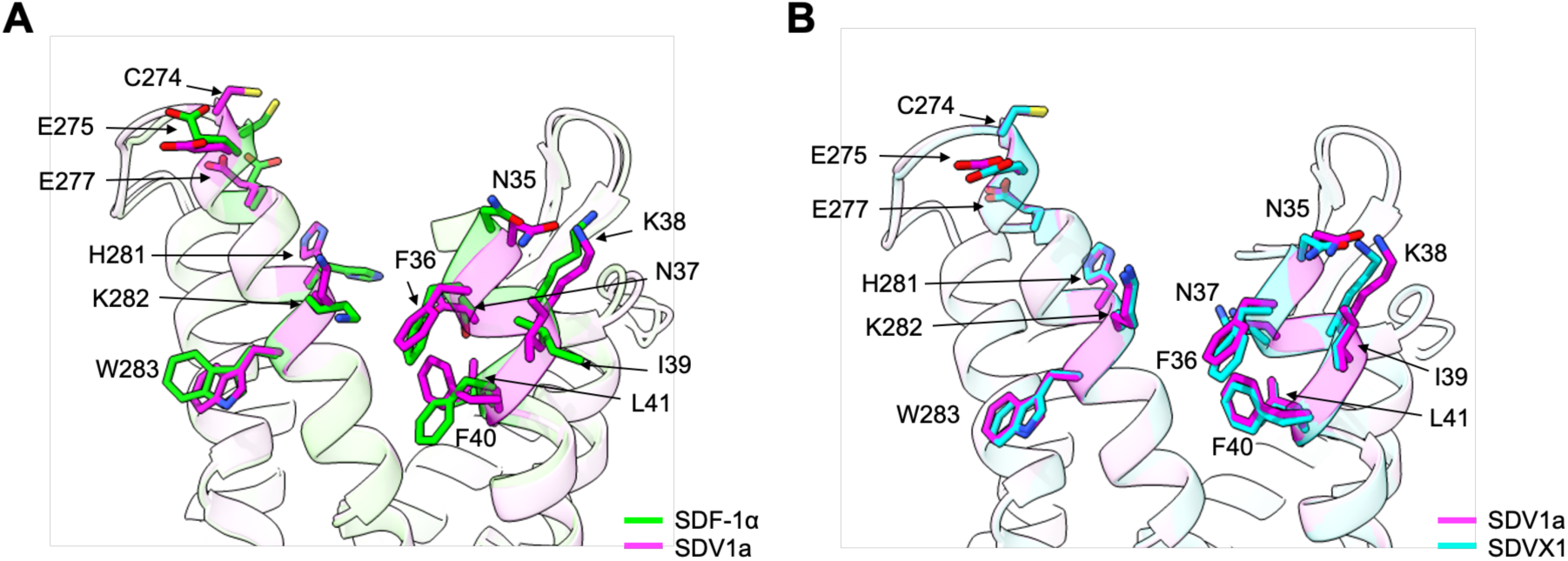
Structural comparison of CXCR4 agonist-bound states induced by SDF-1α, SDV1a, and SDVX1. **(A)** Structural variations in TM7 and TM1 between SDF-1α (green) and SDV1a (magenta) binding systems. **(B)** Structural consistency in TM7 and TM1 between SDV1a (magenta) and SDVX1 (cyan) binding systems.

Given that the N-terminal region of CXCR4 lacks a stable secondary structure, we were concerned that the observed residue perturbations of TM1 might result from the intrinsic structural instability of CXCR4 rather than ligand binding. To address this concern, we further compared the effects of SDV1a and SDVX1 on the above mentioned TM1 residues. The results indicated that the structural changes induced by SDV1a and SDVX1 in this region were highly consistent with no significant perturbations observed (Fig. 5B). Furthermore, ΔRRCS calculations between SDV1a and SDVX1 systems revealed total seven residue pairs (only three residue pairs of CXCR4) with |ΔRRCS| values exceeding 3 (Extended Data Table 3), underscoring the high structural similarity between the two systems. Notably, both SDV1a and SDVX1 share the same LGASWHR binding motif, further supporting the ligand-specific origin of the residue’s conformational perturbations.

### Functional validation of the cryo-EM’s structural information on CXCR4-SDV1a interaction using site-directed mutagenesis

We conducted site-directed mutagenesis analysis to ascertain the functional roles of the residues important for CXCR4-SDV1a signaling as suggested by the cryo-EM structure. In parallel, mutational analysis of CXCR4-SDF-1α signaling was also performed for comparison. Most mutations exhibited similar activation levels for both SDV1a and SDF-1α (Fig. 3D), indicating similar signaling mechanisms which is consistent with the close structural mimicry seen in their cryo-EM structures described above. Mutations such as D193A, W195A, Q200A, D262A, and H281A, which are located in the major pocket and primarily maintain pocket structure, had minimal impact on signaling (Fig. 3D), as they neither strongly interact with the N-terminal amino acids of SDF-1α nor disrupt the critical conformational network for G-protein coupling. In contrast, mutations like Y45A, W94A, D97A, Y116A and E288A, which are situated in the minor pocket and interact with the N-terminal amino acids of SDF-1α and SDV1a, play key roles in receptor activation by stabilizing ligand binding and transmitting structural changes essential for G-protein coupling. These mutations led to significantly reduced signaling (Fig. 3D). Notably, certain mutations, including R30A, D97A, V112A, T117A and I284A, impaired SDV1a signaling more than SDF-1α (Fig. 3D). Overall, these mutagenesis results are consistent with the cryo-EM structures and validate the structural and functional mimicry by our synthetic molecules of CXCR4’s natural agonist ligand.

### Comparative analyses of CXCR4 structures bound by agonists versus antagonists to reveal plausible CXCR4 activation mechanisms

To unveil the structural mechanisms of CXCR4 activation, we systematically compared three agonist-bound structures (SDF-1α ^21^, SDV1a, and SDVX1 reported here) with six previously resolved antagonist-bound structures (IT1t ^13^, CVX15 ^13^, vMIP-II ^14^, HF51116 ^15^, AMD070 ^15^ and AMD3100 ^15^), conducting pairwise analyses across all 18 agonist-antagonist structure comparison pairs. RRCS analysis provided important insights into CXCR4 activation mechanism in several aspects: (1) Comparison of the SDF-1α-bound CXCR4 structure with six antagonist-bound structures revealed notable RRCS differences across eleven residue pairs, forming a continuous signaling pathway that spans three functional regions on CXCR4: the ligand-binding pocket (D97^2.63^-R183^ECL2^, H113^3.29^-R188^ECL2^, L41^1.35^-Y45^1.39^, Y45^1.39^-W94^2.60^, Y45^1.39^-E288^7.39^), the transmembrane signal transmission domain (W252^6.48^-Y255^6.51^, W252^6.48^-A291^7.42^, L127^3.43^-F248^6.44^, S123^3.38^-H294^7.45^), and the G protein binding region (R77^3.29^-Y302^7.53^, R134^3.50^-Y219^5.58^) (Extended Data Table 4). These findings highlight the interconnected nature of conformational changes caused by ligand binding, linking extracellular ligand-binding events to G protein recruitment at the intracellular side. (2) Analysis of the two synthetic agonist SDV1a- and SDVX1-bound CXCR4 structures compared to the six antagonist-bound structures revealed that eight of the residue pairs mentioned above still exhibit notable RRCS differences, consistent with the SDF-1α-bound system, though overall ΔRRCS values are diminished (Extended Data Table 4). These observations suggest that SDV1a and SDVX1 activate CXCR4 in a manner similar to the natural agonist SDF-1α, but through a diminished and less extensive signaling pathway, potentially reflecting their differences in CXCR4 activation efficiency. (3) RRCS analysis of CXCR4 structures bound by AMD070 and AMD3100, both classified as antagonists, revealed activation-like RRCS (ΔRRCS_actives-AMD070/AMD3100_ close to zero) in two residue pairs—Y45^1.39^-W94^2.60^ and W252^6.48^-A291^7.42^ (Extended Data Table 4). These residue pairs, close to the ligand-binding region, indicate that AMD070 and AMD3100 can induce localized conformational changes at these sites but fail to propagate downstream signaling required for full receptor activation. This finding aligns with their reported partial agonist activity ^23^, suggesting that their structural impact on CXCR4 is insufficient to trigger a complete receptor signaling cascade chain.

Further structural comparison of residues contributing to significant ΔRRCS across active and inactive states of CXCR4 revealed significant conformational changes in eight key residues: W94^2.60^, E288^7.39^, F248^6.44^, W252^6.48^, Y302^7.53^, R134^3.50^, R77^2.43^ and Y219^5.58^ (Fig. 6A). These residues were consistently implicated across the comparisons, highlighting their critical roles in CXCR4 activation and agonism. These identified residues align closely with conserved motifs and structural elements reported in GPCR activation studies. For example, the TM6 transmission switch (F248^6.44^, W252^6.48^) is consistent with the hallmark outward movement of TM6 observed in GPCR activation ^22,24–26^, which facilitates G protein recruitment. Similarly, the tyrosine toggle switch (Y219^5.58^ and Y302^7.53^) and the ionic lock residue (R134^3.50^) have been implicated as critical microswitches in GPCR signaling, coordinating conformational changes across the transmembrane helices ^27–29^. These identified eight residues are distributed across essential functional regions of CXCR4, with W94^2.60^ and E288^7.39^ in the ligand-binding pocket identified as the initial switches that trigger activation signals. These signals are subsequently propagated through the TM6 transmission switch (F248^6.44^, W252^6.48^), to the G protein binding region (the ionic lock R134^3.50^ in TM3 and the tyrosine toggle switch Y219^5.58^ and Y302^7.53^ in TM5 and TM7, respectively) (Fig. 6A). Together, these residues form a continuous signaling propagation pathway that connects ligand-binding events at the extracellular pocket to G protein recruitment at the intracellular side.

**Figure 6.**
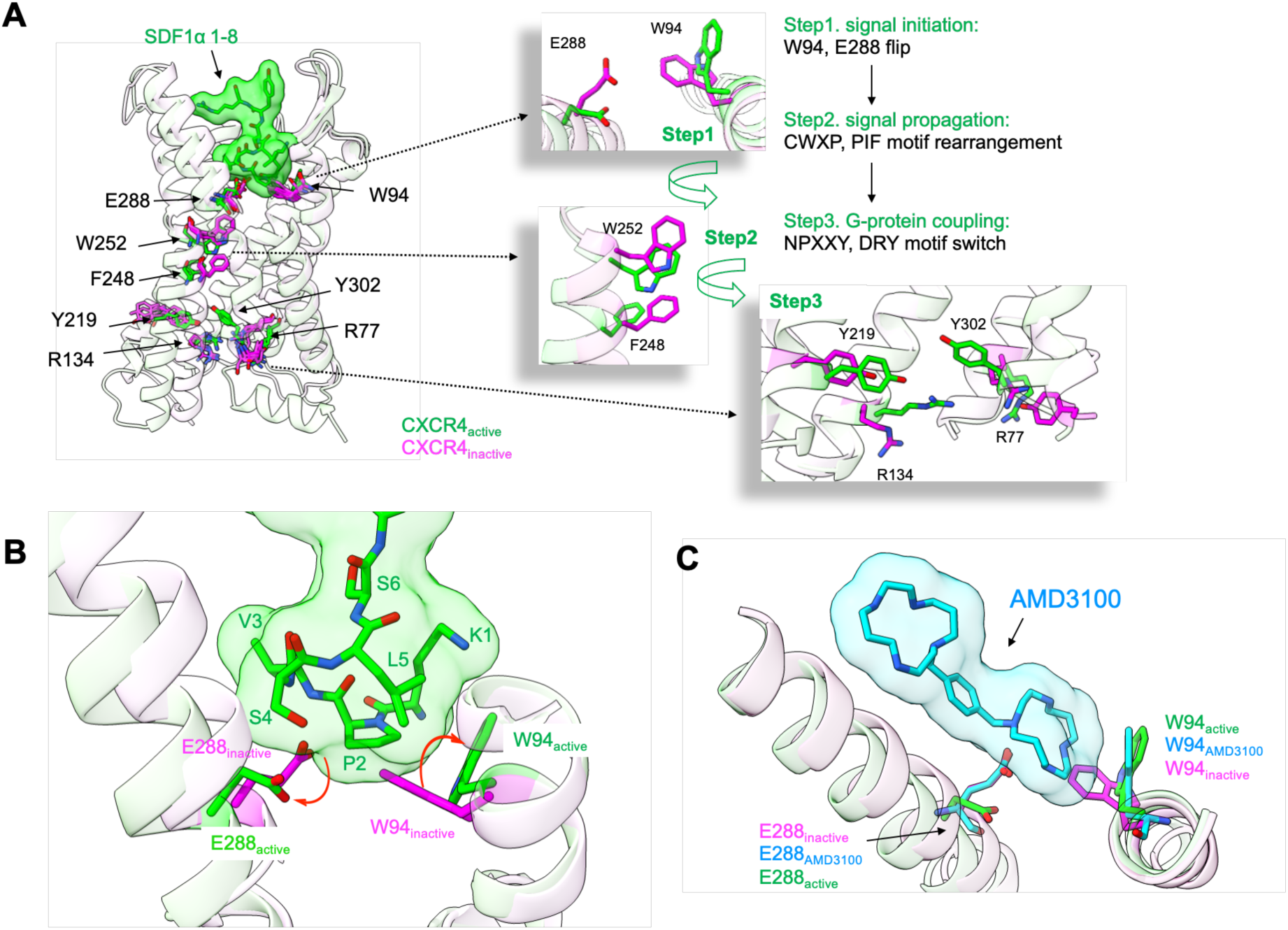
A plausible model for the structural mechanism of CXCR4 activation by its agonist ligand and synthetic mimic molecules. A group of eight residues were identified by comparative analyses of agonist-versus antagonist-bound CXCR4 structures to form a potential signal propagation pathway that relays the effects of conformational changes of key residues such as W94 and E288 caused by extracellular ligand binding to residues downstream involved in G protein recruitment. **(A)** Eight residues (W94^2.60^, E288^7.39^, F248^6.44^, W252^6.48^, Y302^7.53^, R134^3.50^, R77^2.43^, and Y219^5.58^) showing large conformational changes from the inactive (magenta: IT1t, CVX15, vMIP-II, HF51116) to active states (green: SDF-1α, SDV1a, and SDVX1) and their roles in the three-step activation model. **(B)** Conformational changes of W94^2.60^ and E288^7.39^ triggered by the N-terminal eight residues of SDF-1α. **(C)** Conformational changes of W94^2.60^ and E288^7.39^ induced by AMD3100. These results from comparative structural analyses highlight the potential critical roles of eight CXCR4 residues for receptor activation upon agonist interactions, particularly among them W94 and E288 seemingly serving as the initiating sites. They provide consistent structural explanation for not only the structural and functional mimicry of agonist SDF-1α by its synthetic mimics SDV1a and SDVX1, but also the partial agonist effect of antagonist drug AMD3100.

Previous studies on GPCR activation mechanisms suggested that signal transmission begins at the CWXP motif (W252^6.48^) ^24,25,30^. However, the CWXP motif is relatively distant from the ligand-binding pocket, leaving unclear how agonist-induced conformational changes in the pocket propagate to the CWXP motif. Our newly resolved agonist-bound CXCR4-Gi complex structures here reveal that W94^2.60^ in the ligand-binding pocket adopts a consistent vertical conformation in the active state, contrasting sharply with its parallel conformation in the antagonist-bound state ^15^. Additionally, the side chain of E288^7.39^ exhibits distinct conformations between the active and inactive states (Fig. 6B). Notably, AMD3100 and AMD070, despite being classified as antagonists, exhibit atypical structural features that partly align with activation-associated conformations. Specifically, W94^2.60^ in both structures adopt the same conformations observed in the active state. AMD3100, in particular, displays a vertical W94^2.60^ conformation not only in our resolved inactive-state structure but also in the active-state structure reported by others ^21^. However, while AMD3100 and AMD070 activate W94^2.60^, the conformation of E288^7.39^ remains consistent with the antagonist-bound state (Fig. 6C). This structural divergence may explain why AMD3100 has been reported as a partial agonist in previous studies ^23^.

Interestingly, the vertical conformation of W94^2.60^ in the active state is not unique to CXCR4 but has also been observed in other chemokine receptor structures. In the cryo-EM structures of atypical chemokine receptor 3 (ACKR3) activated by SDF-1α, a similar vertical orientation of the equivalent conserved tryptophan residue (W100^2.60^) was reported ^31^. Likewise, in CCR5 structures activated by chemokines such as MIP-1α, the conserved tryptophan (W86^2.60^) rotates vertically which is considered essential for CCR5 activation ^32^. However, its role has not been extensively discussed in the context of the recently resolved SDF-1α-bound CXCR4-Gi cryo-EM structures ^21,33^. This omission underscores the need for further investigation into how W94^2.60^ conformational transition may influence receptor activation and downstream signaling, particularly its relationship with other key residues such as E288^7.39^. Together, these observations suggest that W94^2.60^ and E288^7.39^ may serve as initial signaling sites for CXCR4 activation, while their conserved roles observed across chemokine receptors suggest possible broader implication for general GPCR activation mechanisms.

### Translational benefit of synthetic CXCR4 agonism for stem cell-mediated neuroprotection in a neurodegeneration model

Diseases that are characterized by widespread pathology require a therapeutic intervention that is similarly expansive. Historically in the neural stem cell (NSC) field, mouse models of lysosomal storage diseases (LSDs) have served as prototypes of cerebral diseases that require therapeutic enzyme replacement throughout the brain ^34–37^. The degree of recovery is matched by the extent of enzyme replacement by NSCs secreting these metabolic housekeeping enzymes in the vicinity of enzyme-deficient mutant host cells which they cross-correct. One such disease, Sandhoff Disease (SD), is caused by a deletion mutation of the genes encoding hexosaminidase (Hex) A and B. Previously, we observed that human NSCs (hNSCs), which are intrinsically migratory and constitutively secrete Hex A/B, when implanted in the brains of SD mice, still had a limited reach, leaving areas of host brain uncorrected. However, when SDV1a was administered to distant brain sites at the same time as hNSC implantation, we achieved prolonged symptom-free SD mouse survival by enabling more host cells to be cross-corrected by Hex A/B-secreting hNSCs over a greater expanse ^18^. It could be argued that cell-based enzyme replacement is not very challenging, in that it does not really tap into the multimodal therapeutic machinery of the stem cell because this action could just as effectively be accomplished using viral vectors like AAV9 that can widely disseminate their payload. In the present study, therefore, we sought to demonstrate that the migration-promoting action of a synthetic CXCR4 agonist was critical for directing the migration of hNSCs that exert their therapeutic impact by multiple actions, many of which require the presence of the hNSCs and cannot be mimicked by a drug or gene therapy. Amyotrophic lateral sclerosis (ALS) – an incurable lethal neurodegenerative disease characterized by the inexorable death of α-motor neurons (MNs) in the ventral horn (VH) of the spinal cord (SC) – represents such a disorder.

Previously we reported ^38^ that transplanted undifferentiated, multipotent, NSCs could (in a dose-response manner) reliably forestall the onset-of-disease and substantially prolong symptom-free survival in a large subset of ALS mice (the high-copy SOD1^G93A^ transgenic mouse ^17^) when an implantation technique was employed that ensured chimerism of broad expanses of SC, particularly regions mediating life-sustaining functions (e.g., respiratory control in the cervical SC, coordinated hindlimb function for righting, ambulating, grooming & maintaining an upright posture to feed in the lumbar SC; truncal tone in the thoracic region). Beneficial actions included preservation of diseased host MNs which resulted in a delay-in-symptom onset, preservation of motor function, and prolonged lifespan reliably in 25% of mice. The NSC derivatives that mediated this neuroprotection were the NSC-derived *non*-neuronal cells which invoked a number actions: they produced multiple neuroprotective and neurotrophic cytokines and induced the production of such cytokines in *host* astrocytes; they altered the fate of mutant *host* NSCs (inhibiting them and their NSC-derived astroglial precursors from elaborating disease-permissive, toxin-producing mutant host astrocytes and rather inducing a shift in their developmental program towards the production of neuroprotective gray matter oligodendrocytes; the donor NSCs themselves replaced mutant host NSCs and astroglial precursors with “benign” NSC-derived properly-functioning trophic “fetal-like” astrocytes and gray matter oligodendrocytes. In their role as “fetal” astrocytes, the NSCs created an anti-inflammatory milieu (including blunting microgliosis), mitigated intraneuronal pathology (e.g., protein aggregates, tangled neurofilaments, axonal debris) by restoring proteostasis, and scavenged free radicals and excitotoxins. Donor NSCs did not repress mutant SOD1^G93A^ expression, but rather its clinical manifestations. Because it was the *phenotype*, rather than the genotype, that was influenced, this modulatory approach seemed applicable to a range of ALS etiologies in an accessible, near-term manner.

However, a *critical observation* was that the NSCs’ beneficial impact was directly associated with the degree of diseased neuroaxis they covered: an increase in the number of SOD1^G93A^ SC sites rendered chimeric with normal NSCs was associated with a 30% decrease in *risk-of-death ratio* (Hazard Rate [HR]) (HR=0.70, 95% CI: 0.62-0.79] ^38^. Stated another way, although the response to the NSCs was dose-dependent, the *expanse* of SC terrain covered by the NSCs was more critical than simply increasing the number of NSCs. And, although a broader distribution of donor NSCs across the SC was associated with greater inhibition of pathological progression, it was not only the *extent* of coverage, but ensuring that the terrain rendered chimeric *included regions that subserved life-sustaining functions*; functional improvement was linked to anatomical *location* of the NSC-host chimerism. This expanse of chimerism, however, was achieved by invasively (laminectomies, prolonged anesthesia, blood loss) implanting the NSCs *intraspinally* – into the *VH* and *central canal* at the 4 cardinal SC levels (L_1_,L_3_,T_10_,C_6_). Using that approach, disease *onset* and *progression* were slowed such that *survival for ∼1 year* was reliably achieved in 15-25% of high-copy SOD1^G93A^ mice (untreated control survival = 120-160 days), with some animals surviving *525 days*. Doubling survival was typically achievable in up to 40%. (Equivalent survival in human ALS patients has yet to be achieved). Lifespan was correlated with the number of electrophysiologically-measured functional motor units preserved in the host ^38^.

However, while 25% of mice fared well, 75% were less remarkable – and all ultimately succumbed. *Achieving better coverage* (i.e., replicating what worked in those 25% in the other 75%), and re-dosing the mice with “fresh hNSCs” when they begin to deteriorate, would be at the expense of being even more invasive (i.e., more intraspinal injections “tattooing” the spinal cord and repeating that trauma recurrently). The motivation to circumvent these unacceptable approaches gave rise to testing the adjunctive use of these synthetic CXCR4 agonists with NSCs in the spinal cord (Fig. 7).

**Figure 7.**
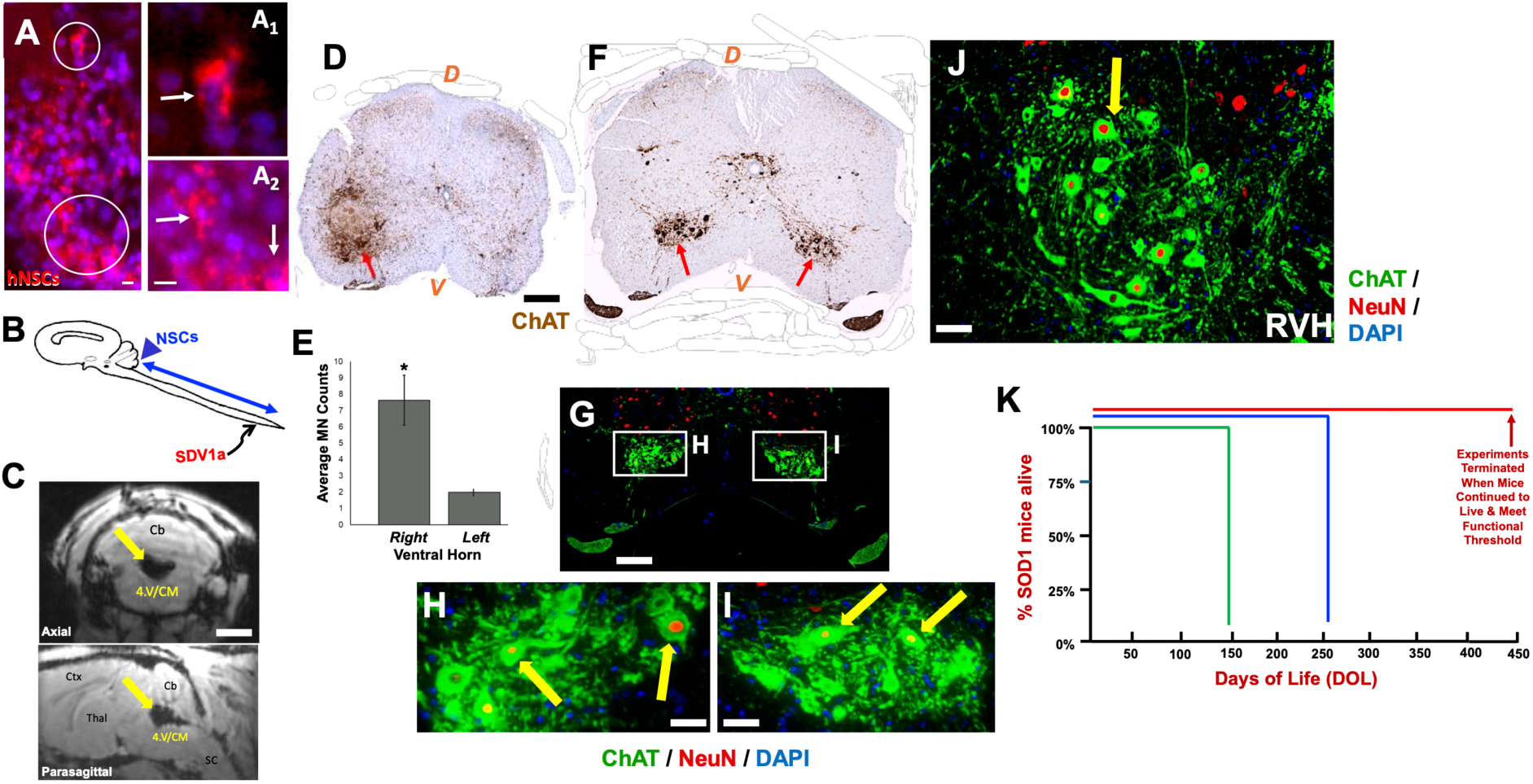
Use of CXCR4 synthetic agonist SDV1a to enhance the spread of neuroprotective hNSCs along the diseased spinal cord (SC) of the SOD1^G93A^ mouse model of ALS (while also minimizing the invasiveness of treatment and retreatment) leads to increased survival of ventral horn MNs and prolongation of symptom-free survival. **(A)** hNSCs and SDV1a in the SC parenchyma were compatible. Representative SOD1^G93A^ mouse with stably engrafted & dispersed hNSCs (detected by STEM 121; human cytoplasm marker; red) following SDV1a injection, visualized 1 month following intraparenchymal transplant of hNSCs showing tissue incorporation with host (**non-red**) cells. 2 representative regions (**circled**) are magnified in **A1 & A2**. SDV1a augmented the survival & migrational reach of the hNSCs. **(B)** Schematic of less invasive administration of an hNSC suspension into the *cisterna magna (CM)* rostrally **(blue arrowhead)** with co-administration of SDV1a into the lumbar spinal cord parenchyma caudally **(arrow)**. **(C)** Magnetic Resonance Image (MRI) in real time of a representative living SOD1^G93A^ mouse having received a successfully placed injection into the CM of hNSCs (pre-labeled with superparamagnetic iron oxide to enable visualization) The dark signal void in axial & parasagittal slices represents the hNSCs in proper location **(yellow arrows)**. Abbreviations: 4.V: 4th ventricle, Cb: cerebellum, Ctx: cortex, Thal: thalamus, SC: spinal cord. **(D)** Histological section through the lumbar SC of an SOD1^G93A^ mouse, stained by immunoperoxidase for ChAT to identify ventral horn MNs, that received an injection of SDV1a injection to only the right side and, hence, drew neuroprotective hNSCs to predominantly that side. A significant number of **ChAT+** MNs **(red arrow)** (quantified in **E**, **\***p<0.01) are preserved on the treated right side as opposed to a dearth of MNs on the uninjected left side, serving as an internal control of the putative efficacy of both the chemoattraction of the SV1a and the neuroprotective action of the hNSCs in the same mutant mouse. **(ChAT+** cells were counted in **E** only if they are also **NeuN+** to increase specificity and rigor). **(F-H)** Immunohistochemistry of a section through the lumbar SC of a representative long-lived (440 days) SOD1^G93A^ mouse who received retreatment with hNSCs and SDV1a bilaterally when motor symptoms became apparent. [F] A cluster of ChAT**+** α-MNs in the ventral horn are visualized at low power using immunoperoxidase **(red arrows)** and in **(G-I)** using dual immunofluorescence to demonstrate that the ChAT+ cells **(green)** co-stain for the mature neuronal perinuclear marker NeuN **(red)** identified by **yellow arrows** in **H** and **I** which are magnification of the boxed areas in **G**, respectively. **(J)** Magnified view of the rescued ChAT**+**/NeuN**+** cells **(yellow arrow)** in the L5 ventral horn of a long-lived SOD1^G93A^ mouse that had been re-dosed with hNSCs + SDV1a and is represented as one of the mice in the **red curve** of the Kaplan-Meier curve in **K**. **(K)** Kaplan-Meier Survival Curve of SOD1^G93A^ mice that received hNSCs into the CM & SDV1a into lumbar SC. Mutants treated with hNSCs alone into the CM (**blue**) (n=2) survived longer (250 days) than untreated mutants (**green**) (n=50). Survival was enhanced in mutants that received a second course of hNSCs into the CM + paired with SDV1A into the lumbar SC at the first sign of symptoms (∼240 DOL) (loss of BBB points) (**red**) (n=3); this cohort was sacrificed at 440 DOL not because they were failing (they were still alive and meeting locomotor criteria) but to confirm that their survival was due to MN neuroprotection, confirmed by histology and quantification in **D-J**. **Abbreviations:** D = dorsal; V = ventral; ChAT = choline acetyl transferase (a marker for a-MNs); NeuN = Neuron nuclei (a marker for mature neurons); **RVH** = right ventral horn. **Scale bar** = 400 µm in **D** & **F**; 20 µm in **G-I**; 30 µm in **J**.

The most non-invasive way to administer cells to the human SC is intrathecally. The most accessible rostral ingress to the thecal sac, from which CSF circulates down and around the SC, is into the *cisterna magna (CM)*, which is quite approachable in patients via a thin percutaneous needle using landmarks or ultrasonic guidance. No surgical procedure is required. It is also known that the NSCs administered into the CSF space will enter the ventral horn by “crawling” along the ventral roots ^39^. We reasoned that the powerful chemoattraction of these peptide agonists for hNSCs such as SDV1a would enhance the migration of hNSCs in the intrathecal space to the caudal end where the SDV1a chemoattractant peptide would be injected minimally invasively into parenchyma (a modification of a lumbar puncture), having already established that SDV1a and the hNSCs are quite compatible with each other in the SC (Fig. 7A and 7B). MNs in the SOD1 mice typically degenerate symmetrically, caudal-to-rostral.

First, we determined the baseline efficacy of hNSCs alone via the CM route-of-administration (ROA). We treated high-copy SOD1^G93A^ mutant mice on 60-70 days-of-life (DOL) (just prior to symptom onset, as is routine in the ALS field) with an intra-CM infusion of allogenic human NSCs (hNSCs) in suspension (n=5) ^38^. Targeting was accurate as illustrated by real-time MRI (Fig. 7C). To prevent immunorejection of hNSCs, all mice – experimental and controls – received oral tacrolimus starting 3 days pre-infusion and continuing until sacrifice or death; the immunosuppressive drug alone was previously established to have no effect on disease onset or progression ^38^.

Mice injected with hNSCs alone lived and met motor criteria (rotarod, Basso-Beattie-Bresnahan (BBB) and Tarlov scales) more than 3 months longer than untreated natural history controls for that colony (Fig. 7K, blue curve vs. green curve). Because the intra-CM ROA is less invasive and traumatic than the prior intraparenchymal ROAs that we and others have published which require laminectomies ^38^, we re-treated a small cohort of mice at the first sign of functional decline (20% reduction in rotarod or BBB/Tarlov scoring) (∼240 DOL) with another course of hNSCs into the CM, but this time adding the SDV1a chemoattractant (1.28 mM in 2 μL) at the opposite end of the SC – into the lumbar (L5) SC parenchyma via non-traumatic, non-surgical intervertebral insertion of an ultrathin (33 gauge) needle – to enhance the SC coverage by the hNSCs. Those animals continued to live with minimal symptoms, far beyond the natural history of the disease. In fact, they were living to at least 440 DOL at which time we terminated the experiment to perform MN counts hypothesizing that the MNs must have been protected to explain the functional integrity and long lifespan of the mice. The mice were healthy and would have continued to live much longer had we not sacrificed them to perform SC histology. SDV1a alone does not alter natural history.

The histology was striking (Fig. 7D-J). MNs are classically identified by immunostaining for the enzyme choline acetyl transferase (ChAT). Proof-of-concept was demonstrated in dramatic fashion in the representative mouse shown in Fig. 7D, which received its SDV1a injection asymmetrically into only the right SC leaving the untreated left SC as an internal control. On the left side of the histological section of that mouse’s lumbar SC, which did not receive SDV1a, few if any ChAT+ MNs are apparent, as one might expect in an untreated mutant SOD1^G93A^ mouse. However, on the animal’s right side which *did* receive SDV1a and drew neuroprotective hNSCs to that region, we see the survival of host MNs (Fig. 7D, red arrow), presumably helping to preserve the motor function of such long-lived high-copy SOD1^G93A^ mutant mice (Fig. 7K, red curve). Fig. 7E shows quantification of this putative SDV1a-mediated asymmetry in MN survival.

In the majority of animals in which both sides of the SC received SDV1a, ChAT+ MNs were protected bilaterally (Fig. 7F-H). To reiterate, mutant SOD1^G93A^ copy number remained high in these long-lived ALS mice; loss of copy number did not account for their longer survival. Indeed, the representative mice shown in Fig. 7D-J had copy numbers >20 and as high as 30.

We know hNSCs that do not integrate long-term into the parenchyma via this ROA; so there was no expectation for them to be detectable more than a half-year post-transplant. In no case was there cell overgrowth, tumors, cell types inappropriate to the CNS, or deformation of normal cytoarchitecture. In no treated animal was lifespan, function, pain, alertness, or comfort level worse than in control mice.

Taken together, these findings suggest that SDV1a is effective in enhancing the spread of neuroprotective hNSCs. These experiments further suggested that SOD1^G93A^ mice – and ALS patients – likely require re-dosing with a neuroprotective regime (in our case hNSCs + SDV1a) to optimize survival. Re-treatment at the first sign of disease re-progression can be lifesaving. The animals so treated here lived for nearly a year longer than untreated mice, and at least a half-year longer than mice treated just once. Indeed, they would have lived longer had the experiment not been terminated for histological assessment of MN presence given no signs of functional decline.

## Discussion

While *antagonism* of the chemokine receptor CXCR4 has been extensively studied and approved for clinical use, selective CXCR4 *agonism* by synthetic molecules is an unanticipated therapeutic intervention we recently unveiled ^18^. The structural mechanisms which enable them to perform therapeutic CXCR4 signal activation, however, are poorly understood but could help elucidate a new class of drugs. Here in this study, we determined the cryo-EM structures of CXCR4-Gi complexes bound and activated by two de novo designed and chemically synthesized peptides – SDV1a and SDVX1 – dual-moiety molecules that, in a novel fashion, link agonistic moiety with antagonistic moiety to selectively trigger CXCR4’s desirable function in mediating cell migration. These data represent the first reported CXCR4 structures in states activated by synthetic agonist peptides.

It is noteworthy that these synthetic agonists induce conformational changes in CXCR4 that align closely with the structural changes triggered by the natural agonist SDF-1α. This structural consistency supports the notion that the first 1-8 residues of SDV1a and SDVX1 are sufficient to mimic the activation mechanism of full-length SDF1α and effectively drive CXCR4 signaling. In addition, while sequence variations between SDV1a/SDVX1 and SDF-1α impact binding affinity and induce localized conformational differences near the extracellular ends of TM1 and TM7, these changes do not compromise the signaling function of the 1-8 residue segment. This observation highlights the functional independence of the two moieties in our de novo designed peptides: the binding moiety facilitates receptor engagement whereas the chemokine’s N-terminal residues as the signaling moiety initiate receptor signal transmission without mutual interference. Together, these findings validate the dual moiety strategy as an effective approach for CXCR4 agonist design, demonstrating the effectiveness of integrating distinct binding and signaling functionalities within a single ligand. By leveraging the *dual moiety concept*, this approach simplifies the ligand structure while retaining the essential functional elements required for receptor activation, paving the way for the development of smaller and more efficient CXCR4 agonists. The structural and functional mimicry of natural ligand-induced CXCR4 agonism with much smaller and simplified peptides supports this de novo design strategy for other GPCR systems. The translational value of these simplified de novo dual moiety GPCR agonists was demonstrated in our experiments on the SOD1^G93A^ mouse model of ALS. The agonist peptide SDV1a enhanced the multimodal neuroprotection of hNSCs in the SOD1^G93A^ mouse spinal cord by optimizing their spread across the diseased neuroaxis (it having been previously demonstrated by us that the success of neuroprotective actions of transplanted NSCs is directly correlated with the expanse of diseased neuroaxis traversed by the donor cells ^38^). Spinal cord regions that received chemoattractant SDV1a, were those regions where MNs were best protected by the hNSCs to an extent sufficient to preserve locomotor function that enabled survival longer than previously reported for a high-copy mutant. In facilitating this broader neuroprotection, these agonist peptides also enabled two highly impactful logistical changes in the cell-based treatment of ALS. First, they permitted the minimally invasive intrathecal route of administration (via the cisterna magna). Second, they enabled what this study highlighted so dramatically: that SOD1^G93A^ mice – and ALS patients – likely require re-dosing with a neuroprotective regime at the first sign of disease progression to prolong survival. For re-dosing to be practical, it must be easy to re-administer and atraumatic; SDV1a made repeating minimally-invasive intra-CM administration possible, or even contemplating increasing total cell dosage (if cell suspension volume is too great) by simply dividing the administration period over the course of a few days.

Being able to control the migration and distribution of therapeutic stem cells may find applications beyond ALS to other neurodegenerative and neurotraumatic conditions. In conclusion, we report the first high resolution cryo-EM structures of a GPCR, chemokine receptor CXCR4 bound and activated by synthetic peptides mimicking the agonistic action of natural CXCR4 ligand SDF-1α. These two CXCR4 structures in the activated state in complex with G protein and two peptide agonists SDV1a and SDVX1, respectively, provide detailed structural information, not reported before, about the structural mechanism of CXCR4 signaling activated by synthetic molecules. This fills an important gap in the mechanistic understanding of CXCR4 agonism since previous studies mainly focus on CXCR4 antagonism with small molecules including two clinical antagonist drugs. Furthermore, it provides a structural foundation for developing a new class of CXCR4 agonist drugs for combinational therapies with human neural stem cells of ALS and other neurodegenerative diseases.

## Methods

### Peptide synthesis

The peptides SDV1a, SDVX1 were synthesized through the general protocol of solid-phase peptide synthesis, as described previously ^18^. However, unlike SDV1a, SDVX1 was obtained via repeated cycles (Fmoc deprotection step and Amino acid coupling step) of coupling reaction with 20 Fmoc-protected amino acids (Fmoc-AAs), whereas SDV1a required coupling reaction of 31 Fmoc-AAs. The purity of SDVX1 was verified by HPLC (high performance liquid chromatography), while the molecular weight of SDVX1 was analyzed by MALDI-TOF-MS (matrix-assisted laser desorption ionization time-of-flight mass spectrometry).

### CXCR4 binding assay

The CXCR4 binding assay was performed as described previously ^15^. CHO cells stably expressing CXCR4 were seeded into 96-well plates with 5×10^5^ cells/well, followed by adding various concentrations of peptides and mouse anti-CXCR4 antibody 12G5 (BD Biosciences, 555971) in a final volume of 100 μl of assay buffer (PBS with 0.5% BSA and 0.05% NaN_3_). After incubation on ice for 40 min, cells were washed with assay buffer by centrifugation and incubated with anti-mouse IgG-FITC antibody (Sigma-Aldrich, F9137) for 30 min on ice, then washed with assay buffer two times. Finally, the fluorescence intensity was measured by a PerkinElmer EnVision multimode plate reader with an excitation wavelength of 485 nm and an emission wavelength of 535 nm. Experiments were performed in at least two independent replicates. Data are presented as mean ± standard deviation (SD). The IC_50_ values were calculated using GraphPad Prism software.

### CCR5 binding assay

The CCR5 binding assay was conducted following the same protocol as the CXCR4 binding assay, with the following modifications: CHO cells stably expressing CCR5 were used instead of CXCR4-expressing cells, and mouse anti-CCR5 antibody 45523 (R&D Systems, MAB181) was used in place of the CXCR4-specific antibody.

### Cell migration assay

SupT1 cells (2×10^6^ cells/well in 75 μl) were added into the upper chamber of the Transwell-96 well plates with 5.0 μm pore size polycarbonate membrane (Corning, 3387). Then, 200 μl of medium containing 2 nM SDF-1α or various concentrations of peptides was added to the lower chamber. The plates were incubated at 37 ℃ for 3 hours. Migrated cells in the lower chamber were measured using Celltiter-96 reagent (Promega, G3580). To investigate the cell migration promoting mechanism of our two agonists and the natural ligand SDF-1α, cells in the upper chamber were pretreated with the PKC inhibitor Go6983, the PI3K inhibitor Pilaralisib or the ERK1/2 inhibitor FR180204-either alone or in combination - for 15 min. Experiments were performed in at least two independent replicates. Data are presented as mean ± standard deviation (SD).

### Luciferase reporter assays

pGL4.29[*luc2P*/CRE] vector (Promega, E8471), pGL4.30[*luc2P*/NFAT-RE] vector (Promega, E8481), pGL4.33[*luc2P*/SRE] vector (Promega, E1340) or pGL4.34[*luc2P*/SRF-RE] vector (Promega, E1350) was transiently transfected into CHO cells stably expressing CXCR4. After 8 hours of transfection, cells were trypsinized and seeded into 96-well plates at 2×10^4^ cells/well, followed by overnight incubation. Subsequently, cells were serum-starved for 5 hours and then stimulated with various concentrations of agonists. For the CRE luciferase reporter assay, cells were additionally treated with 2.5 μg/mL forskolin to enhance cAMP signaling. After 6 hours of incubation at 37 °C, the culture medium was removed, and 20 μL of cell lysis buffer was added to each well. Then, 100 μL of luciferase assay reagent was added, and luminescence was measured using a microplate reader. Experiments were performed in at least two independent replicates. Data are presented as mean ± standard deviation (SD).

### Site-directed mutagenesis assay for functional analysis of CXCR4

CXCR4 mutants were generated by PCR-based site-directed mutagenesis as described previously ^40^. Plasmids encoding wild-type/mutated CXCR4 were co-transfected with pGL4.33[*luc2P*/SRE] into CHO cells, followed by stimulation with SDV1a (5 µM) or SDF-1α (50 nM), according to the SRE luciferase reporter assay protocol described above.

### Constructs design

Human CXCR4 with N119S mutation was cloned into pFastBac 1 vector with a HA signal sequence followed by a Flag tag. The protein Bril and protein LgBit were fused to the N-terminus and C-terminus of CXCR4 to enhance the CXCR4 expression level and stabilize the CXCR4-Gi complex, respectively. Domain negative (DN) Gαi1 was cloned into pFastBac 1 vector and Gβ1γ2 with HiBit fusion to the C-terminus of Gβ1 was cloned into pFastBac Dual vector. GPCR/G-protein complex-stabilizing antibody scFv16 was also cloned into pFastBac 1 vector containing a N-terminal GP67 secretion signal peptide and a C-terminal 8×His tag.

### Expression of Bril-CXCR4-LgBit, Gαi1, Gβ1γ2 and scFv16 proteins

High-titer recombinant baculovirus (>10^8^ viral particles per mL) was obtained using the Bac-to-Bac Baculovirus Expression System (Invitrogen) as described previously ^15^. Bril-CXCR4-LgBit, Gαi1 and Gβ1γ2 (virus ratio of 2:1:1) were co-expressed in Sf9 insect cells at a density of 2.5×10^6^ cells/mL. Cells were harvested after incubation at 27 °C for 60 hours. scFv16 was expressed in Sf9 insect cells at 27 °C for 72 hours.

### Purification of CXCR4-Gi-scFv16-agonist complex

The cell pellets expressing CXCR4-Gi were suspended and homogenized in a solubilization buffer containing 20mM HEPES, pH 7.5, 300 mM NaCl, 1% LMNG/0.2% CHS, 10% glycerol, 5 mM MgCl_2_, 25 mU/mL apyrase, 50 μM agonist and cOmplete EDTA-free protease inhibitor cocktail for 2 hours at 4 °C. The supernatant was isolated by centrifugation at 18,000 rpm for 30 min, and incubated with anti-DYKDDDDK G1 affinity resin (GenScript, L00432) for 2 hours at 4 °C. After binding, the resin was washed with ten column volumes of 20 mM HEPES, pH 7.5, 100 mM NaCl, 0.1%LMNG/0.02%CHS, 10% glycerol, 5 mM MgCl_2_ and 50 μM agonist, followed by four column volumes of 20 mM HEPES, pH 7.5, 100 mM NaCl, 0.01%LMNG/0.002%CHS, 10% glycerol, 5 mM MgCl_2_ and 50 μM agonist. The protein was then eluted with three column volumes of 20 mM HEPES, pH 7.5, 100 mM NaCl, 0.01%LMNG/0.002%CHS, 10% glycerol, 5 mM MgCl_2_, 50 μM agonist, 0.25 mg/mL DYKDDDDK peptide and cOmplete EDTA-free protease inhibitor cocktail. The protein was concentrated using Amicon Ultra 100 kDa molecular weight cut-off spin concentrator (Millipore, UFC910024) and incubated with scFv16 for 30 min on ice with a molar ratio of 1:1.5. The CXCR4-Gi-scFv16-agonist complex was further purified by size exclusion chromatography on Superose 6 Increase column in buffer 20 mM HEPES, pH 7.5, 100 mM NaCl, 0.01%LMNG/0.002%CHS, 5 mM MgCl_2_ and 50 μM agonist. The peak fraction was collected and concentrated to around 5.0 mg/mL for cryo-EM sample preparation.

### Cryo-EM sample preparation, data collection and data processing

The purified SDV1a-CXCR4-Gi-scFv16 and SDVX1-CXCR4-Gi-scFv16 complexes were deposited onto glow-discharged grids (Zhenjiang Lehua Technology, CryoMatrix M024-Au300-R12/13) and subsequently vitrified using a Vitrobot Mark IV. Images were then acquired in the counted-Nanoprobe mode with a 300 kV Titan Krios Gi3 electron microscope (Thermo Fisher Scientific), which was equipped with a Gatan K3 Summit detector and a GIF Quantum energy filter. Video stacks, consisting of 50 frames each, were collected using SerialEM software in counting mode, set to a nominal magnification of ×105,000, with a pixel size of 0.85 Å and a defocus range of −1.2 to −1.8 μm. Each video stack was recorded for 2.2 seconds, with an exposure time of 0.044 s per frame, resulting in a total dose of 54.2 e−/Å².

For the SDV1a-CXCR4-Gi-scFv16 complex, a total of 3,231 movies were collected and imported into RELION 5.0, where motion correction was conducted using MotionCorr2. The micrographs were then transferred to CryoSPARC, where the CTF estimation was performed using PatchCTF. 6,452,608 particles were auto-picked and extracted in a pixel size of 1.7 Å. Two rounds of 2D classification were performed, and the remaining 1,991,213 particles were used for ab-initio model reconstruction and heterogeneous refinement. The retained 756,448 particles were re-extracted in RELION in a pixel size of 0.85 Å. After 3D refinement in RELION, one round of particle sieving was conducted using CryoSIEVE, resulting in the retention of 211,499 particles. These particles were then subjected to CTF refinement and particle polishing in RELION, as well as non-uniform and local refinement in CryoSPARC. This comprehensive process yielded an electron density map with an overall resolution of 2.8 Å.

For the SDVX1-CXCR4-Gi-scFv16 complex, a total of 6,687 movies were collected and imported into RELION 5.0, where motion correction was conducted using MotionCorr2. The micrographs were then transferred to CryoSPARC, where the CTF estimation was performed using PatchCTF. 11,47,200 particles were auto-picked and extracted in a pixel size of 1.7 Å. Two rounds of 2D classification were performed, and the remaining 3,459,212 particles were used for ab-initio model reconstruction and heterogeneous refinement. The retained 742,993 particles were re-extracted in RELION in a pixel size of 0.85 Å. After 3D refinement in RELION, one round of particle sieving was conducted using CryoSIEVE, resulting in the retention of 194,772 particles. These particles were then subjected to CTF refinement and particle polishing in RELION, as well as non-uniform and local refinement in CryoSPARC. This comprehensive process yielded an electron density map with an overall resolution of 2.7 Å.

### Atomic model building and refinement

The coordinates for CXCR4 from PDB entry 8U4O, along with the coordinates for the N-terminus of SDV1a/SDVX1, Gi, and scFv16 from PDB entry 8HNK, were docked into the density maps using ChimeraX. The resulting complex structure was manually adjusted in Coot and subsequently refined using Phenix. Model geometries were validated with MolProbity, and the structural figures were created in ChimeraX.

### Human Neural Stem Cells (hNSCs)

Human neural stem cells (hNSCs) were obtained, at low passage, from the stable well-characterized cell line HFB2050 ^18,38^. For expansion, hNSCs were thawed and seeded on T-75 flasks coated with Poly-L-Ornithine and Laminin substrates to ensure attachment and post-thaw viability. The cells were then cultured with biweekly media changes of Neurobasal Medium (containing B27 without Vitamin A, 1% Penicillin/Streptomycin, and 1% GlutaMax) supplemented with Normocin, Heparin Sulfate, Leukemia Inhibitory Factor (LIF), and Basic Fibroblastic Growth Factor (bFGF) until 90% confluence was reached, at which point hNSCs were dissociated using Accutase and passaged into tissue culture-treated flasks for further expansion to achieve desired doses for animal studies. For administration to mice, hNSCs were dissociated using Accutase and resuspended at a concentration of 2 x 10^5^ cells/μL in sterile Hank’s Balanced Salt Solution (HBSS).

### hNSC administration to SOD1^G93A^ mouse model of ALS

Pre-symptomatic SOD1^G93A^ mice at 60-70 DOL were randomly selected to receive initial hNSC infusions vs. HBSS alone into the cisterna magna (CM).

It should be noted that intra-CM injections in mice require a surgical procedure whereas the administration in human patients is non-surgical, minimally-invasive, and performed without general anesthesia; cells are simply infused via a thin needle placed percutaneously into the capacious CM, guided by landmarks or ultrasound.

For the mice, 3 days prior to hNSC administration, all animals were started on daily oral Tacrolimus to prevent rejection of the xenograft. Animals were anesthetized with gaseous isoflurane. The dorsal hair at the base of the skull was sheared. Skin was disinfected with ethanol and povidone iodine. The animal was placed on a sterile field and had its head positioned on the Stoelting Stereotactic Injector with constant anesthesia provided as inhaled isoflurane gas. A 25.4-mm sagittal incision was made in a rostro-caudal direction spanning the occipital bone to the first cervical vertebra. The skin and muscles were separated via blunt dissection, followed by retraction. The muscles were cut to expose the CM injection site. An opening to access the subdural space was made by gently piercing the dura mater with a non-invasive 30-gauge needle affixed to hemostat forceps to allow for unobstructed access for the hNSC-loaded syringe. hNSCs were loaded into a Hamilton syringe affixed with 30 gauge needle, stereotactically-positioned, and administered at a controlled rate of 2 μL/minute. These initial doses were between 2.5-5.0 x 10^5^ cells. Dose escalation studies were performed (n = 16) such the ultimate dose of 1 x 10^6^ hNSCs was used for re-treatment. Increasing the dose to 1 x 10^7^ hNSCs mandated a cell suspension volume ≥10µL which caused reflux of the injectate, suggesting that if a greater number of hNSCs was required for effective neuroprotection, the total dose would need to be divided between 2 administrations separated by a few days. The outcome data in Fig. 6K, however, suggests that 1 x 10^6^ hNSCs per infusion should be sufficient for adequate neuroprotection if accompanied by administration of SDV1a.

Upon completion of injection, muscles were sutured with an Ethilon 6-0 suture. Skin was rejoined with 4-0 Vicryl suture. Buprenorphine analgesic and Lactated Ringers were subcutaneously administered and mice were placed back in their home cage for recovery. The animals remained on chronic immunosuppression using oral tacrolimus until death or euthanasia.

When rotarod, BBB or Tarlov scale criteria worsened by 20%, or if a cohort animal precipitously reached euthanasia threshold based on functional decline, a subset of animals, received re-treatment with hNSCs (1 x 10^6^) via the CM injection site previously used for the first infusion. That site was re-exposed using the same technique described above.

Immediately following hNSC administration, SDV1a was injected into the lumbar spine parenchyma. While for the mouse, an intra-spinal injection entails the surgical procedure described below, for human patients, such administration is a non-surgical, minimally invasive technique which is similar to a lumbar puncture.

For the mouse, to expose the lumbar spine, the animal’s dorsal hair was sheared and skin was disinfected with ethanol followed by povidone iodine. To access the lumbar enlargement, a 50.8-mm sagittal incision on the skin made caudo-rostrally. Skin and muscle were separated via blunt dissection. The muscles were gently teased apart to expose the ventral roots. 1.28 mM SDV1a was injected into the spinal cord parenchyma in a 2 μL volume at an upper and lower lumbar site using a non-invasive 30 gauge needle. SDV1a in the lumbar SC parenchyma was not volume limited. After the procedure was completed, animals were administered Buprenorphine analgesic and Lactated Ringers subcutaneously and placed in home cages for recovery.

### Locomotor Testing

Animals were weighed biweekly and were administered locomotor tests weekly. As others have ^38,39^, we adapted the Basso-Bresnahan-Beatty (BBB) scale ^41^ for hindlimb function in SOD1 mice. The BBB scale is a 21-point scale with specifically defined criteria for determining levels of hindlimb function, where “0” and “21” indicate no hindlimb function to normal gait & function respectively. Additionally, we used the Basso Mouse Scale (BMS) which is a mouse-specific translation of the BBB scale to further assess hindlimb function in mice ^42^. We adapted the Wrathall-modified Tarlov scale, originally designed for hindlimb function, to assess forelimb ^43^. Tarlov is a 5-point scale where “0” indicates no forelimb movement and “5” indicates normal forelimb movement and function. The BBB, BMS, and Wrathall-modified Tarlov scales were administered to mice in an open-field setting with two independent and blinded evaluators assigning scores based on the criteria in the scales.

To assess gait, hindlimb & forelimb coordination, core muscle strength, and balance we used the Rotarod. In line with our previous reports ^18,37,38^, animals were deemed to have “passed” the rotarod if they could maintain balance and ambulate on the apparatus moving at a fixed speed of 8 rpm for at least 20 seconds. We determined the time to cut-off the trial at 210 seconds (3.5 minutes). When an animal failed the Rotarod trial (i.e., not remaining on the Rotarod for at least 20 seconds), the animal was administered the righting reflex to determine essential animal neurological function. The mouse was placed in a supine position, and if it failed to right itself within 10 seconds, the animal was euthanized with isoflurane gas.

### Immunohistochemistry

Either at the conclusion of the study or when animals reached euthanasia threshold (inability to right from a supine position within 10 seconds), mice were transcardially perfused with heparinized saline followed by 4% paraformaldehyde to fix the tissue. The entire nervous system was dissected first by progressive laminectomies to expose the dorsal surface of the spinal cord. Transverse processes were gently dissected to allow for spinal cord roots to be left intact. After the dorsolateral spinal cord was exposed, the brain was dissected by gently cutting the skull along the suture joints. Finally, to take out the whole brain and spinal cord, the vertebra body was detached from the spinal cord. The lumbar enlargement was then isolated, cut into 2-mm pieces, and then formalin-fixed paraffin-embedded where 5 μm-thick slices were generated in a caudo-rostral manner.

Following standard deparaffination and rehydration, tissue sections were stained sequentially first using anti-ChAT (1:2000) (to recognize choline acetyl transferase, a well-acknowledged α-MNs marker) followed by NeuN antibodies (Abcam; 1:500) (NeuN being a perinuclear marker that identifies mature neurons). The tyramide signal amplification system (TSA; ThermoFisher) was used. Slides were stained on a Ventana Discovery Ultra (Ventana Medical Systems, Tucson, AZ, USA). Antigen retrieval was performed using a basic treatment solution (CC1; pH 8.5, Ventana) for 40 minutes at 95 °C followed by a peroxidase quenching step with Discovery Inhibitor (Ventana) for 12 min. Sections then were incubated in anti-ChAT antibody for 32 minutes at 37 °C, followed by anti-rabbit-HRP (OmniMap Detection Kit, Ventana) and detected using TSA-Alexa 488. The antibodies were stripped from their epitopes by treatment in a citric acid-based solution, pH 6 (CC2, Ventana) for 24 minutes at 95 °C. Subsequently, the slides were incubated with anti-NeuN antibody for 32 minutes at 37 °C followed by anti-rabbit-HRP (OmniMap Detection Kit, Ventana) and detected using TSA-Alexa 594. Nuclei were labeled with DAPI, and slides were mounted with fluorescent mounting medium (ProLong^TM^ Gold antifade reagent). To amplify the ChAT signal, the HQ-Amp system (Ventana) was used with a 4 min amplification time. After amplification, the anti-HQ-HRP antibody was incubated on the slides followed by development with DAB and a counterstain of hematoxylin. Slides were rinsed, dehydrated through alcohol and xylene and coverslipped.

To calculate the number of α-MNs (as per **Fig. 6E**), ChAT**+** cells were counted in 3 representative sections 20 μm apart through a given spinal segment using unbiased stereology. An apparently ChAT**+** cell was confirmed to be a neuron by observing its co-expression of NeuN.

## Data availability

The cryo-EM density maps for the SDV1a-CXCR4-Gi-scFv16 and SDVX1-CXCR4-Gi-scFv16 complexes have been deposited in the Electron Microscopy Data Bank under accession numbers EMD-64402 and EMD-64403, respectively. The corresponding atomic coordinates for SDV1a-CXCR4-Gi-scFv16 and SDVX1-CXCR4-Gi-scFv16 have been deposited in the Protein Data Bank under accession number 9UPU and 9UPV, respectively. All other data are included in the manuscript and/or the supplementary information.

## Acknowledgments

This work was supported by the California Institute for Regenerative Medicine (DISC2-12666), the NIH (GM057761), the Ciechanover Institute of Precision and Regenerative Medicine, the Shenzhen-Hong Kong Cooperation Zone for Technology and Innovation (HZQB-KCZYB-2020056), and the Tsinghua-Peking Center for Life Sciences. This work was also supported by the National Natural Science Foundation of China (Project No. 32100963 and 32471254, to H.H.), Shenzhen Science and Technology Innovation Committee (Project No. JCYJ20210324131802008 to H.H.), the Kobilka Institute of Innovative Drug Discovery, Presidential Fellowship and University Development Fund at the Chinese University of Hong Kong, Shenzhen (H.H. and H.J.), Shenzhen Outgoing Postdocs Research Fund (J10120230692 to H.J.), and Shenzhen Medical Research Fund, Youth Project (A2403008 to H.J.). This work was also supported in part by grants to E.Y.S. from the Department of the Army Medical Research (HT94252510320), the Martha M. Mack Foundation, and the Richmond Family Foundation for Research into Pediatric Brain Disorders. A.C. is supported by grants from Adelson Medical Research Foundation (AMRD), the Israel Science Foundation (ISF), the Israeli Personalized Medicine Partnership (IPMP), and a Professorship from the Israel Cancer Research Fund (ICRF) USA. We thank the Kobilka Cryo-EM Center at the Chinese University of Hong Kong, Shenzhen, for supporting EM data collection.

## Author contributions

Z.H., J.A. and E.Y.S. conceived of the project and oversaw its overall execution; H.H. oversaw structural determination; E.Y.S. oversaw the animal studies; X.S. produced the protein; H.J. conducted the structural determination; Q.M. performed the *in vitro* studies; X.F. designed and synthesized the peptide; K.S.S., M.A.L, L.V.Z., A.I.W.A., R.L.N., J.O., V.O., Y.X., I.S.H. participated in the animal studies and related microscopy and quantification; J.Z. performed structural comparison analysis; D.P.P performed histology; Z.H., Y.X., X.S., H.J., H.H., Q.M., X.F., X.L., Y.M., T.Q. A.C., J.A., E.Y.S. interpreted results; Z.H., E.Y.S., H.H., X.S., H.J., J.Z, J.A. wrote the manuscript.

## Additional information

**Supplementary Information** is available for this paper.

**Correspondence & requests for Materials** should be addressed to Ziwei Huang, Hongli Hu, Evan Y. Snyder, or Jing An.

## Extended Data Figures and Tables

**Extended Data Figure 1.**
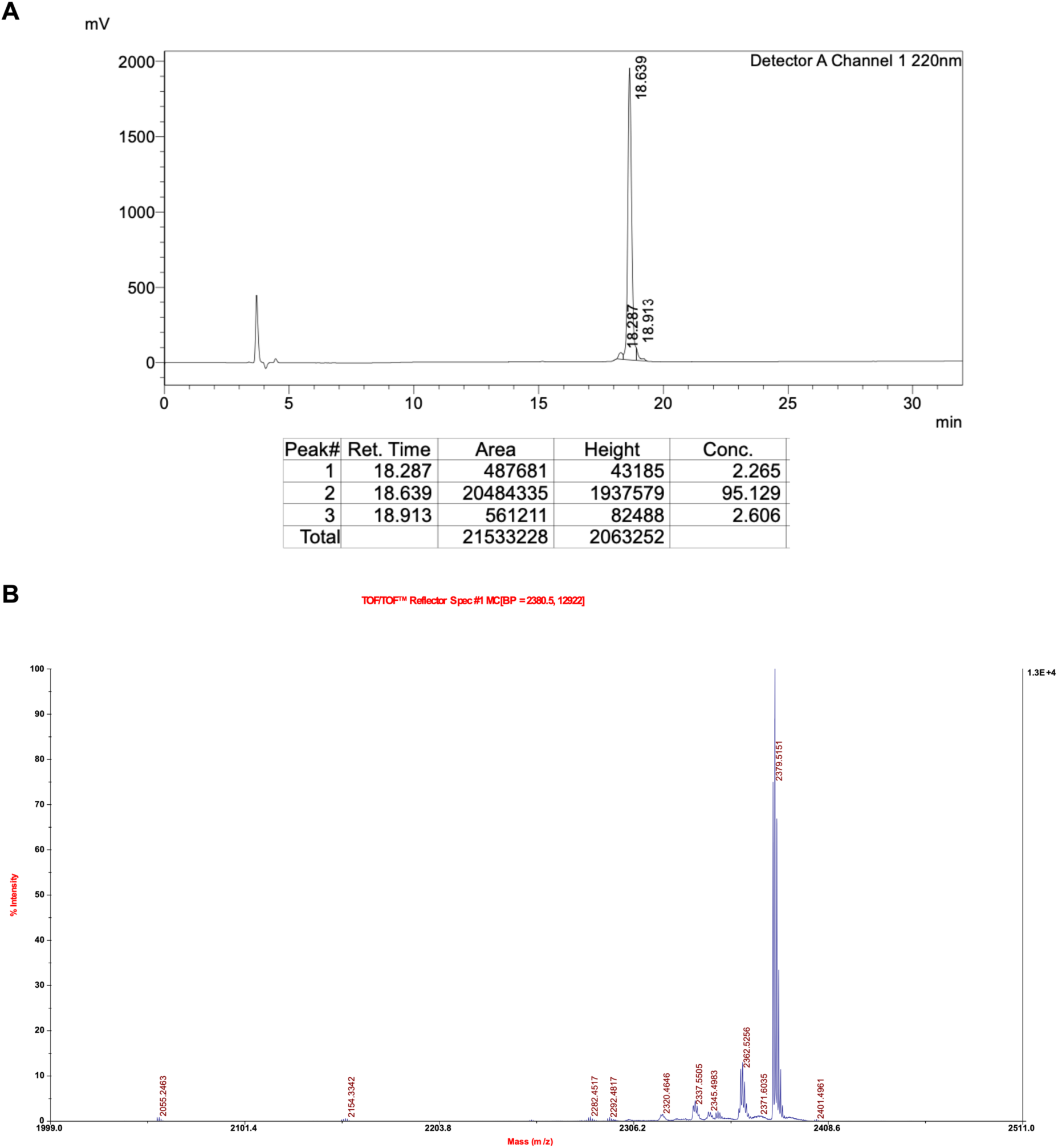
Validation of purity and molecular weight of SDVX1. (A) The purity of SDVX1 was checked by HPLC (high performance liquid chromatography). (**B**) The molecular weight of SDVX1 was analyzed by MALDI-TOF-MS (matrix-assisted laser desorption ionization time-of-flight mass spectrometry).

**Extended Data Figure 2.**
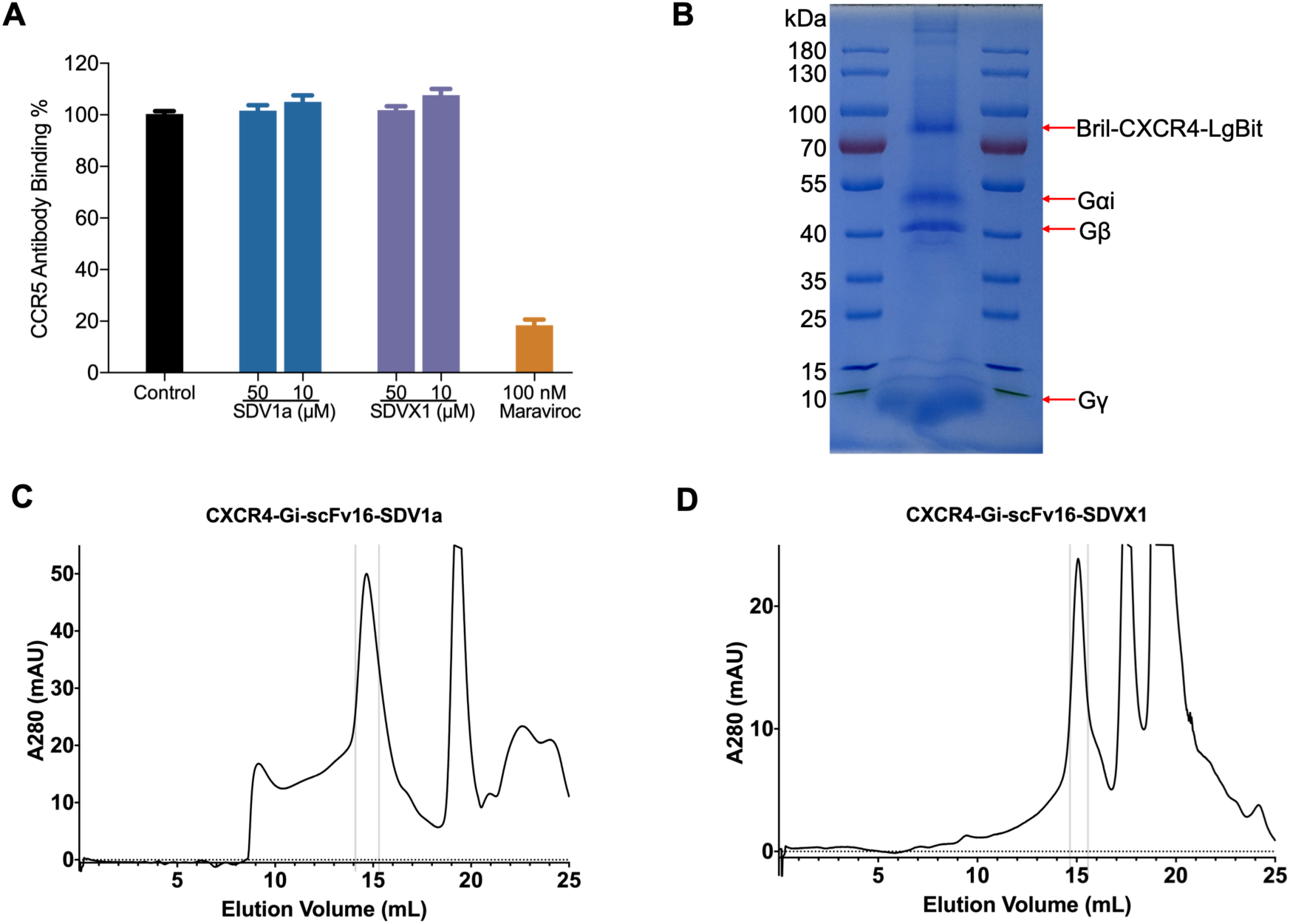
CCR5 competitive binding assay and purification of CXCR4 complexes. **(A)** The binding affinity of SDV1a and SDVX1 on CCR5 receptor. Maraviroc as a positive control. **(B)** Coomassie-stained SDS-PAGE gel of purified CXCR4-Gi complex. **(C)** The elution curve of the SDV1a-CXCR4-Gi-scFv16 complex on a Superose 6 Increase column. **(D)** The elution curve of the SDVX1-CXCR4-Gi-scFv16 complex on a Superose 6 Increase column.

**Extended Data Figure 3.**
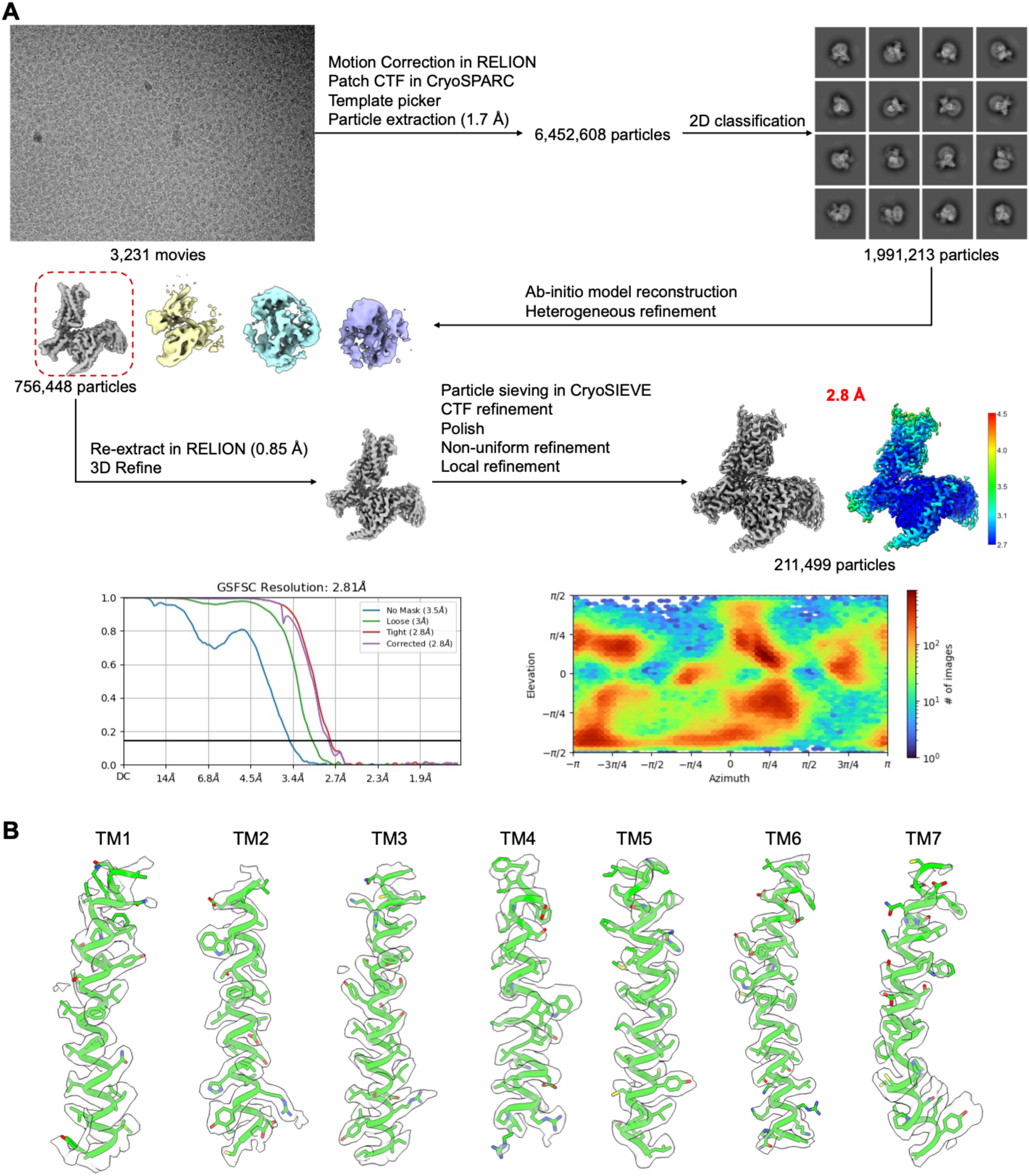
Data processing of SDV1a-CXCR4-DNGi-scFv16 complex. **(A)** Data processing flow of the cryo-EM dataset of the SDV1a-CXCR4-DNGi-scFv16 complex. The local resolution map, the FSC curve, and the statistics of the orientation distribution of the particles are presented in the diagram. **(B)** Local electron densities of the transmembrane helices.

**Extended Data Figure 4.**
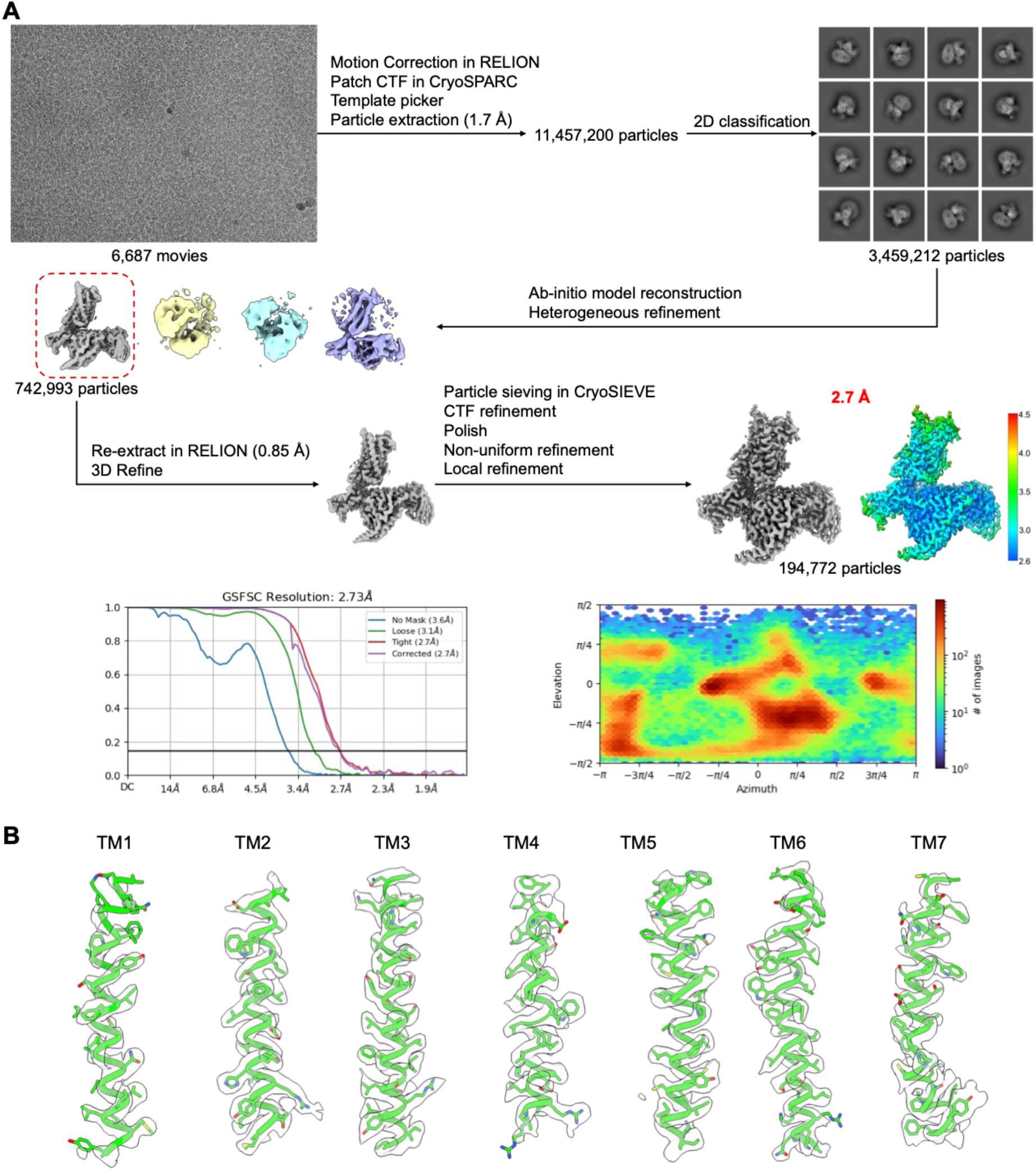
Data processing of SDVX1-CXCR4-DNGi-scFv16 complex. **(A)** Data processing flow of the cryo-EM dataset of the SDVX1-CXCR4-DNGi-scFv16 complex. The local resolution map, the FSC curve, and the statistics of the orientation distribution of the particles are presented in the diagram. **(B)** Local electron densities of the transmembrane helices.

**Extended Data Table 1.** Statics of data collection, data process, model refinement and validation.

|  | **SDV1a-CXCR4-Gi-scFv16**<br>**(EMDB-64402)**<br>**(PDB 9UPU)** | **SDVX1-CXCR4-Gi-scFv16**<br>**(EMDB-64403)**<br>**(PDB 9UPV)** |
| --- | --- | --- |
| **Data and processing** |  |  |
| Magnification | 105,000 | 105,000 |
| Voltage (kV) | 300 | 300 |
| Electron exposure (e<sup>-</sup>/ Å<sup>2</sup>) | 54.2 | 55.2 |
| Defocus range (μm) | -1.2 ~ -1.8 | -1.2 ~ -1.8 |
| Pixel size (Å) | 0.85 | 0.85 |
| Symmetry imposed | C1 | C1 |
| Initial particle images (no.) | 6,452,608 | 11,457,200 |
| Final particle images (no.) | 211,499 | 194,772 |
| Map resolution (Å) | 2.8 | 2.7 |
| FSC threshold | 0.143 | 0.143 |
| Map resolution range | 2.7 ~ 4.5 | 2.6 ~ 4.5 |
| **Refinement** |  |  |
| Initial model used (PDB code) | - | - |
| Model resolution (Å) | 3.0 | 3.0 |
| FSC threshold | 0.5 | 0.5 |
| Map sharpening B factor (Å<sup>2</sup>) | 103.0 | 100.0 |
| Model composition |  |  |
| Non-hydrogen atoms | 8,837 | 8,837 |
| Protein residues | 1,135 | 1,135 |
| Ligands | 1 | 1 |
| B factor (Å<sup>2</sup>) |  |  |
| Protein | 54.50 | 73.95 |
| Ligand | 79.84 | 96.53 |
| R.m.s. deviations |  |  |
| Bond lengths (Å) | 0.002 | 0.002 |
| Bond angles (°) | 0.467 | 0.484 |
| Validation |  |  |
| MolProbity score | 1.28 | 1.31 |
| Clash score | 3.08 | 2.74 |
| Rotamers outliers (%) | 0.11 | 0.00 |
| Ramachandra plot |  |  |
| Favored | 96.96 | 96.25 |
| Allowed | 3.04 | 3.75 |
| Outliers | 0.00 | 0.00 |

**Extended Data Table 2.** Residue pairs with |ΔRRCS| values exceeding 3 identified in CXCR4 systems bound by SDF-1α and SDV1a.

| Residue 1 | Residue 2 | RRCS<br>(SDF1α-CXCR4) | RRCS<br>(SDF1α-CXCR4) | ΔRRCS<br>(SDF1α-SDV1a) |
| --- | --- | --- | --- | --- |
| R:117_THR | R:171_ASP | 0.6 | 5.7 | -5.1 |
| R:190_TYR | R:199_PHE | 2.6 | 7.3 | -4.7 |
| R:176_ASN | R:189_PHE | 1.0 | 5.3 | -4.2 |
| R:232_HIS | R:236_LYS | 0.1 | 3.9 | -3.9 |
| R:35_ASN | R:38_LYS | 0.0 | 3.7 | -3.7 |
| R:79_HIS | R:161_TRP | 0.6 | 4.4 | -3.7 |
| R:40_PHE | R:41_LEU | 0.0 | 3.0 | -3.0 |
| R:223_ILE | R:238_LEU | 0.4 | 3.4 | -3.0 |
| R:122_SER | R:161_TRP | 4.0 | 1.0 | 3.1 |
| R:200_GLN | R:262_ASP | 3.7 | 0.5 | 3.2 |
| R:75_LYS | R:153_GLU | 3.8 | 0.5 | 3.3 |
| R:102_TRP | R:184_TYR | 5.1 | 1.7 | 3.4 |
| J:4_SER | R:281_HIS | 3.5 | 0.0 | 3.5 |
| R:113_HIS | R:171_ASP | 6.5 | 2.9 | 3.6 |
| R:36_PHE | R:282_LYS | 4.0 | 0.0 | 4.0 |
| R:100_ALA | R:183_ARG | 4.6 | 0.4 | 4.1 |
| R:36_PHE | R:40_PHE | 7.5 | 3.3 | 4.1 |
| R:101_ASN | R:103_TYR | 11.5 | 7.1 | 4.4 |
| R:178_SER | R:185_ILE | 7.5 | 3.1 | 4.4 |
| R:230_LYS | R:235_ARG | 6.1 | 0.2 | 5.9 |
| R:41_LEU | R:45_TYR | 8.1 | 1.5 | 6.6 |

**Extended Data Table 3.** Residue pairs with |ΔRRCS| values exceeding 3 identified in CXCR4 systems bound by SDV1a and SDVX1.

| Residue 1 | Residue 2 | RRCS<br>(SDV1a-CXCR4) | RRCS<br>(SDVX1-CXCR4) | $\Delta\text{RRCS}$<br>(SDV1a-SDVX1) |
| --- | --- | --- | --- | --- |
| R:272_GLN | R:276_PHE | 0.9 | 6.6 | -5.6 |
| R:255_TYR | R:288_GLU | 2.9 | 7.9 | -5.0 |
| J:8_ARG | R:262_ASP | 1.9 | 6.8 | -4.9 |
| J:2_PRO | R:288_GLU | 2.7 | 7.4 | -4.7 |
| R:101_ASN | R:183_ARG | 4.2 | 8.5 | -4.3 |
| J:1_LYS | J:6_SER | 4.2 | 0.0 | 4.2 |
| J:4_SER | R:288_GLU | 7.8 | 3.0 | 4.8 |

**Extended Data Table 4.** ΔRRCS analysis across all 18 agonist-antagonist structures.

| Res 1 | Res 2 | ΔRRCS<br>(sdf1α-IT1t) | ΔRRCS<br>(sdf1α-CVX15) | ΔRRCS<br>(sdf1α-<br>vMIPII) | ΔRRCS<br>(sdf1α-51116) | ΔRRCS<br>(sdf1α-070) | ΔRRCS<br>(sdf1α-3100) | ΔRRCS<br>(sdv1a-IT1t) | ΔRRCS<br>(sdv1a-CVX15) | ΔRRCS<br>(sdv1a-<br>vMIPII) | ΔRRCS<br>(sdv1a-51116) | ΔRRCS<br>(sdv1a-070) | ΔRRCS<br>(sdv1a-3100) | ΔRRCS<br>(sdvx1-IT1t) | ΔRRCS<br>(sdvx1-CVX15) | ΔRRCS<br>(sdvx1-vMIPII) | ΔRRCS<br>(sdvx1-51116) | ΔRRCS<br>(sdvx1-070) | ΔRRCS<br>(sdvx1-3100) |
| --- | --- | --- | --- | --- | --- | --- | --- | --- | --- | --- | --- | --- | --- | --- | --- | --- | --- | --- | --- |
| H113 | R188 | -5.4 | -2.7 | -3.1 | -3.8 | -3.7 | -2.9 | -5.4 | -2.7 | -3.1 | -3.8 | -3.7 | -2.9 | -5.1 | -2.4 | -2.8 | -3.5 | -3.4 | -2.6 |
| D97 | R183 | -3.9 | -4.2 | -4.2 | -1.5 | -5.7 | -3.6 | -2.7 | -3.0 | -3.0 | -0.3 | -4.4 | -2.4 | -3.1 | -3.4 | -3.4 | -0.7 | -4.8 | -2.8 |
| Y45 | W94 | -6.2 | -5.8 | -1.7 | -3.3 | -0.4 | 0.0 | -6.5 | -6.1 | -2.0 | -3.6 | -0.7 | -0.3 | -6.5 | -6.0 | -1.9 | -3.5 | -0.7 | -0.3 |
| Y45 | E288 | 2.7 | 2.8 | 2.5 | 2.5 | 2.5 | 2.7 | 1.1 | 1.2 | 0.9 | 0.9 | 0.8 | 1.0 | -0.1 | 0.0 | -0.3 | -0.3 | -0.3 | -0.1 |
| L41 | Y45 | 4.2 | 5.8 | 6.4 | 7.6 | 7.4 | 3.9 | -2.4 | -0.8 | -0.2 | 1.0 | 0.8 | -2.7 | -1.6 | 0.0 | 0.6 | 1.8 | 1.6 | -1.9 |
| W252 | A291 | -3.9 | -3.6 | -3.4 | -2.6 | -0.9 | -1.0 | -3.0 | -2.7 | -2.5 | -1.7 | 0.0 | -0.1 | -3.2 | -2.8 | -2.6 | -1.8 | -0.1 | -0.2 |
| W252 | Y255 | -3.6 | -2.2 | -2.6 | -3.4 | -2.8 | -2.4 | -3.1 | -1.7 | -2.1 | -2.9 | -2.3 | -2.0 | -2.8 | -1.4 | -1.8 | -2.6 | -2.0 | -1.7 |
| L127 | F248 | -3.3 | -2.1 | -2.3 | -1.5 | -2.3 | -2.4 | -3.6 | -2.3 | -2.6 | -1.8 | -2.5 | -2.7 | -3.7 | -2.4 | -2.7 | -1.9 | -2.6 | -2.8 |
| S123 | H294 | 3.1 | 2.7 | 3.3 | 3.0 | 3.3 | 2.9 | 0.5 | 0.1 | 0.7 | 0.4 | 0.7 | -2.1 | 0.4 | 0.0 | 0.6 | 0.3 | 0.6 | 0.2 |
| R77 | Y302 | -14.5 | -9.7 | -6.4 | -4.8 | -5.3 | -4.9 | -15.9 | -11.1 | -7.8 | -6.2 | -6.6 | -6.2 | -16.0 | -11.2 | -7.9 | -6.3 | -6.8 | -6.4 |
| R134 | Y219 | 3.7 | / | / | 3.3 | 3.6 | 3.7 | 3.0 | / | / | 2.6 | 2.9 | 3.0 | 2.9 | / | / | 2.5 | 2.8 | 2.9 |

